# Genomic differentiation of British oaks

**DOI:** 10.64898/2026.03.05.708962

**Authors:** Louise Gathercole, Rômulo Carleial, Nathan Brown, Sandra Denman, Ernest Wu, Gabriele Nocchi, Richard Nichols, Richard Buggs

## Abstract

The morphological continuum between Britain’s two native oak species helped Darwin formulate his “one long argument” in *The Origin of Species*. Here we comprehensively analyse the genome-wide differentiation of 418 individuals from 60 British oak populations. Using 7 million biallelic SNPs, we find evidence for two species with islands of variation containing 8,230 SNPs of which 2,005 are within gene annotations. We find extensive hybridisation and back-crossing between the species, and this is biased in favour of introgression from *Quercus robur* into *Q. petraea*. Chloroplasts are shared between the species, mainly of haplotypes derived from Iberian glacial refugia. Genomic allocation of species and hybrids allows analysis of niche differentiation, showing that a larger *Q. robur* component is disproportionately found in warmer and more thermally variable environments with alkaline soils, while a larger *Q. petraea* component is found in more topographically complex terrain with high rainfall. This leads us to emphasise the balance between the opposing forces of hybridisation and niche differentiation in explaining the continuum between the two species. Using long-term field data, we show that stem growth over three decades was higher in *Q. robur* than in *Q. petraea* or hybrids, but this difference could be explained by environmental factors. In contrast, the five triploid trees in our sample grew significantly faster than diploids even after accounting for environmental effects, suggesting that triploids could be a valuable economic resource.

## Introduction

It has long been acknowledged that native oaks in Britain, and especially Scotland, show continuity of variation between two taxa^1^. Darwin noted in *The Origin of Species* that “in this country the highest botanical authorities and practical men can be quoted to show that the sessile and pedunculated oaks are either good and distinct species or mere varieties”^2^. This observation helped him come to the conclusion that varieties and species form a continuum that arises via descent with modification.

Twentieth century morphologists suggested that the oak continuum was caused by hybridisation and backcrossing^3–5^, stimulating extensive debate^6–10^. Subsequent genetic studies showed sharing of chloroplast haplotypes between species^11^, large numbers of hybrid acorns^12^, and genome-wide introgression^13^ between what are now known as *Quercus robur* L. (pedunculate oak) and *Q. petraea* (Matt.) Liebl. (sessile oak). These species tend to be better demarcated in continental Europe^14^, but even there, intermediate individuals are often difficult to classify^14–18^ and molecular studies show extensive hybridisation^11,12,19–22^. Some botanists still view *Q. petraea* as a subspecies of *Q. robur*^23^, and the two tend to be lumped together in forest inventories and in some legislation^24^.

Palynological evidence, which cannot distinguish between the two species, suggests that oaks expanded northwards in Britain after the last glaciation^7,25,26^. Without differential selection, hybridisation could cause the two species to merge into one. Although distribution maps tend to show both species occurring across most of Britain^27–29^, the geographic distribution of the two species is suggestive of niche adaptation. It has long been reported that *Q. robur* is more abundant to the south and east whilst *Q. petraea* is more frequent in the west and north^5,7,30–32^. This has led to the inference that *Q. petraea* is more acid tolerant than *Q. robur* and can thrive at higher altitudes and in shallow soils^5,30,31,33–35^. There has also been selection by humans, with *Q. robur* the preferred species for planting in Britain^5,34,36,37^. On the continent, there is considerable overlap of the two species’ niches^33,38^. However, *Q. petraea* is more common on slopes, hilltops, and free-draining soils^39–41^, is more drought-tolerant^42,43^ and has higher water use efficiency^39,44^, whereas *Q. robur* is frequently associated with lower-lying and wet sites and seems to be more tolerant to waterlogging^45,46^. In Russia and Scandinavia, *Q. robur* has the more northerly range limit^33^.

In Britain and on the continent, genomic regions have been found that are differentiated between the two species^13,47–50^, though these regions differ among studies. Leaf morphological differences between the two species have been shown to be controlled by several quantitative trait loci^51^. There is some evidence that introgression between the two species can be adaptive^52^. Direction of introgression is also affected by an asymmetrical reproductive barrier: *Q. robur* is fertilised easily by *Q. petraea* whereas *Q. petraea* and hybrids are less easily fertilised by *Q. robur*^53–56^. This asymmetry led to the “resurrection model”, whereby *Q. robur* initially colonises new areas through acorn dispersal with *Q. petraea* invading later via pollen swamping^15^, but empirical support for this hypothesis remains mixed^57,58^.

It is important to understand differentiation and population dynamics of *Q. robur* and *Q. petraea*, to inform our understanding of evolution and because oaks support more than 2,000 associated macro-species and provide valuable timber resources^30,59^. Here we report a large whole genome sequence survey of oaks in Britain. We resequence individually, and at high depth, the whole genome of 418 individuals from 60 natural oak populations in Forest Condition Survey Plots across Great Britain^60–62^. We analyse these data to characterise their ploidy and population structure, their hybridisation and genetic differentiation, and their growth and niche differentiation.

## Results

### Genotyping, SNP discovery and ploidy

We identified 22,178,235 SNPs high quality SNPs after filtering (see Methods). The inbreeding coefficient (F) for all 439 samples using a subset of 24,297 SNPs had a mean of 0.085 (SD = 0.05). Six outlier samples showed comparatively more negative coefficients (Supplementary Figure 1) suggesting high levels of excess heterozygosity. One of these samples was found to be anomalous due to a pipeline error and was excluded. An analysis of allelic balance suggested that the remaining five individuals were triploid as they had a majority of SNPs with the proportion of reads supporting each allele at 0.33/0.67, rather than 0.5/0.5 as is more typically seen in diploid individuals. The suspected triploids were single trees from five locations (Supplementary Figure 2) and constitute 1.2% of the total number of samples. A preliminary fastSTRUCTURE analysis prior to triploid removal (data not shown) suggested that two are *Q. robur*, two are *Q. petraea* and one is a back-crossed admixed individual that was predominantly *Q. petraea*. Suspected triploids were removed from the dataset prior to filtering the loci for allele balance, linkage disequilibrium pruning and filtering for minor allele frequency 0.002 and 10% missingness. The filtered dataset (i.e. **dataset 1**, see Methods) had a final count of 7,112,139 SNPs (Table 1).

**Table 1.**
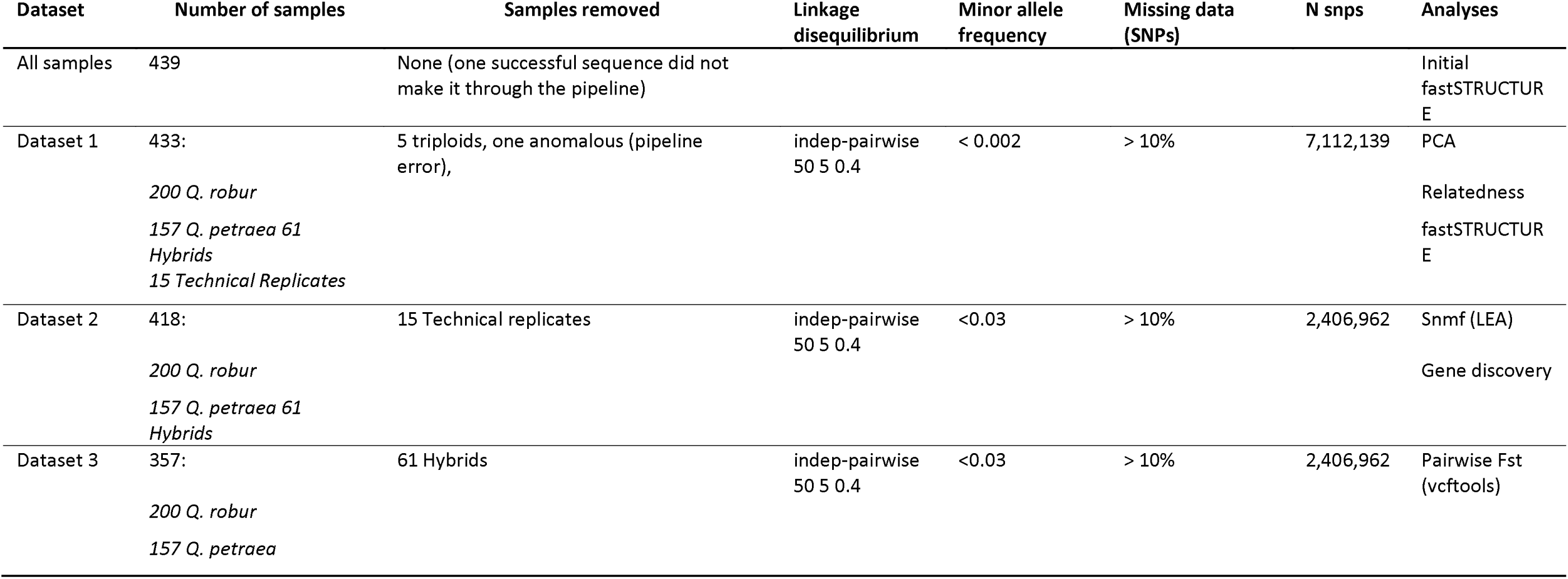
– Summary of SNP datasets used in analyses.

### Population structure and species assignment

The fastSTRUCTURE admixture plots and the function chooseK applied to **dataset 1**, indicated two ancestral clusters (K=2) (Figure 1a). Two clusters were also found in the PCA, in which PC1 clearly separates the samples into *Q. robur* and *Q. petraea*, with an admixed group of hybrids lying in a continuum intermediate to the two species (Supplementary Figure 3). The *Q. petraea* cluster is more dispersed than the *Q. robur* cluster, suggesting greater genetic variation within *Q. petraea*. Both clusters were also identified by snmf analysis.

**Figure 1.**
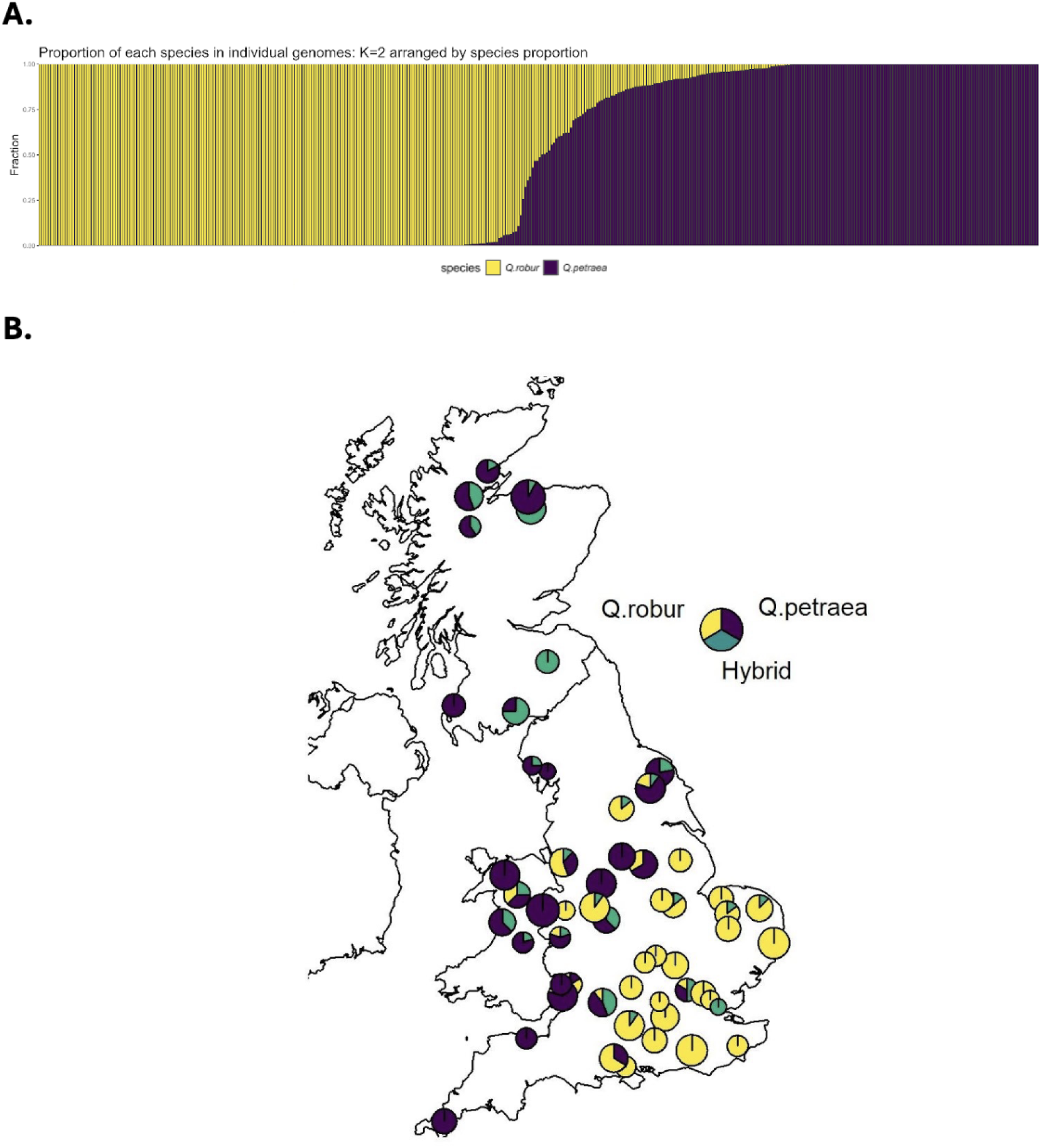
– Population structure in British oak: **(a)** Individual admixture proportions across multiple populations. Each vertical bar is an individual with the proportion of the genome allocated to each species using fastSTRUCTURE, samples arranged by proportion of species 1 (*Q. robur*) using dataset 1 (replicate samples removed). Bars are colour coded by species (gold = *Q. robur* and purple = *Q. petraea*). **(b)** Geographical distribution of two native species of oak across Britain. Each pie chart represents a population (n = 60) and are colour coded according to the proportion of individuals from each species (*Q. robur* = yellow; *Q. petraea* = purple; hybrids = green). Triploid individuals are included in this map.

Using 10% admixture (q) as the cut off for pure versus hybrid status, and excluding the technical replicates from consideration, 200 individuals were assigned to *Q. robur*, 157 to *Q. petraea*, and 61 classified as hybrids/admixed individuals (0.1 < q < 0.9). Considerable allele sharing can be seen and this is asymmetrical, with genetic material being predominantly introgressed from *Q. robur* into *Q. petraea* (Figure 1a). Among the 61 hybrids there were 17 F_1_ hybrids (4%; 0.375 < q < 0.625), 39 samples back-crossed with *Q. petraea* (q < 0.375) and only five back-crossed with *Q. robur* (q > 0.625).

We found that *Q. robur* is predominantly located in the south and east of Great Britain, while *Q. petraea* is mostly in the north and west (Figure 1b). However, there are three sites in the South East of England, where we find *Q. petraea*. Hybrids are mostly found in the north and west of Britain and located in areas where the other sampled trees are mostly *Q. petraea.* At two sites all the samples are hybrids: site 28, an oakwood in the north of Scotland, and site 63 in the borders area of Scotland. We found no pure *Q. robur* samples in the Scottish sites.

Relatedness analysis identified the correct 15 pairs of technical replicates and one extra, unexpected pair. A re-inspection of the labels revealed that the same tree had been processed twice with different IDs, so the mislabelled replicate was removed. After removing replicates, we created a new dataset by filtering SNPs with maf > 0.03, which reduced the SNP set to 2,406,962 SNPs (**Dataset 2,** Table 1).

### Chloroplast haplotype structure

We assembled the full chloroplast sequence (∼160 kb in length) of 151 *Q. petraea*, 61 hybrids, and 178 *Q. robur* individuals and constructed a median joining haplotype network. This has a similar topology to that of Nocchi et al^13^ and we similarly identified node clusters (Figure 2a) in our network that correspond to haplotypes identified by previous restriction enzyme studies^11,63^. Most of our samples were in nodes that correspond to restriction enzyme haplotypes 10, 11, and 12, all containing a diagnostic point mutation in the *trnD*–*trnT* fragment characteristic of chloroplast lineage B. We found two individuals from Hazelborough wood with a highly differentiated haplotype that our network joined by a long branch to a node within haplotype 12. This may correspond to haplotype 1 of lineage C^63^ and we mark it as such in Figure 2, although the matching between our data and previous restriction enzyme gel fragments was not entirely conclusive. We identified nine trees with very divergent chloroplasts that match chloroplast haplotypes 7 or 26 within lineage A^21^ in Gloucester and Chester (Figure 2b). As argued by Nocchi et al^13^, haplotype 7 is the most likely identification as this is widespread in France and central Europe^21^.

**Figure 2.**
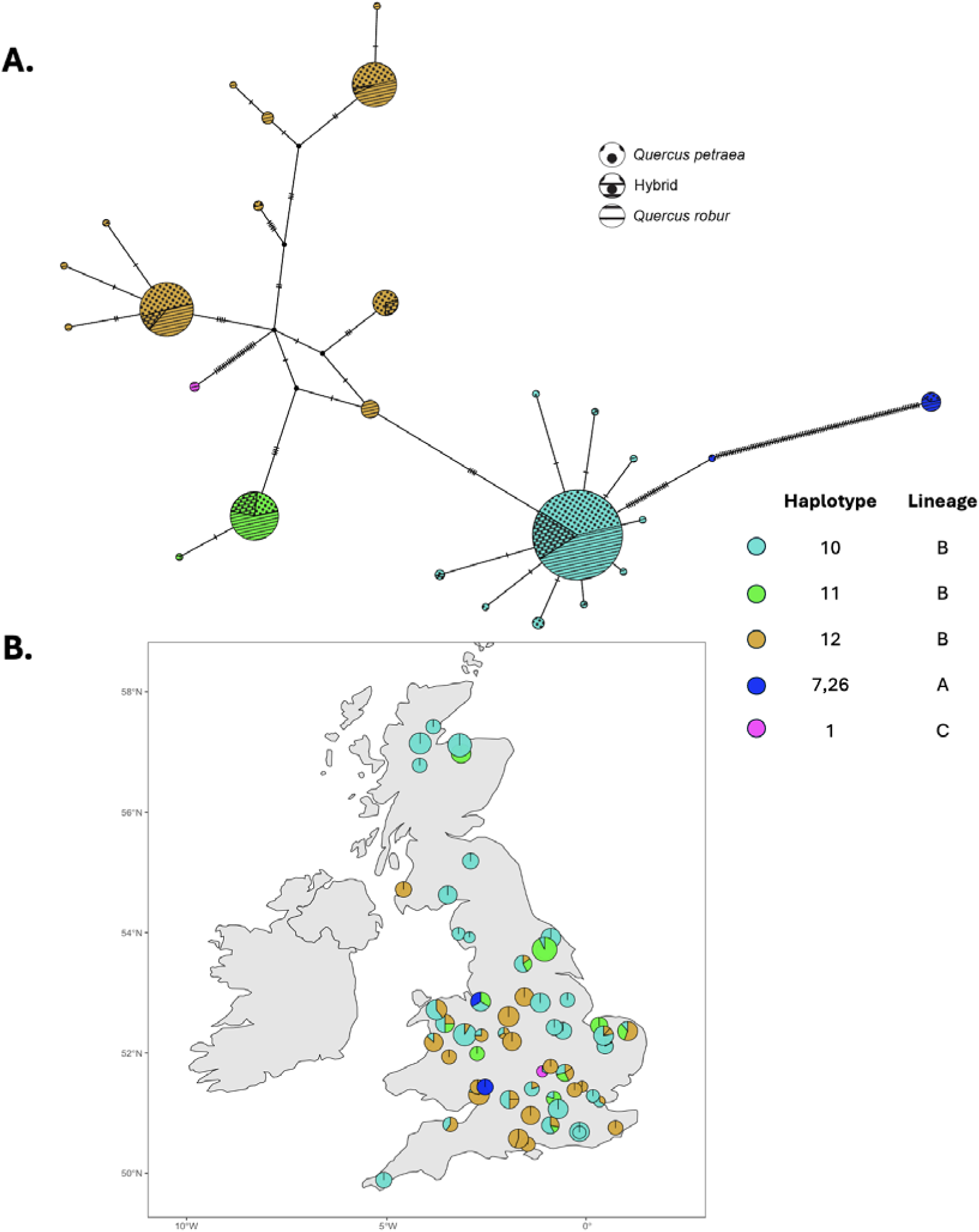
– Chloroplast haplotype network of British oaks: **(a)** Median joining chloroplast haplotype network of 151 *Q. petraea*, 61 hybrids, and 178 *Q. robur* individuals. The nodes are coloured to match the lineage and haplotype identity described in Petit et al. (2002). The sizes of the nodes are proportional to the number of individuals within that haplotype, and the hatch marks along the edges represent the number of single nucleotide mutations between the nodes. Lines and dots within nodes represent the species. **(b)** Geographic distribution of each haplotype across Britain.

### Inter-specific variation and species differentiation

Between *Q. robur* and *Q. petraea* nuclear genomes, the mean interspecific F*_ST_* across SNP loci was 0.03 (median 0.01, standard deviation 0.03, range 0 to 0.98). This was calculated using vcftools between pure *Q. robur* (n=200) and *Q. petraea* (n=157) samples (i.e. **dataset 3)**, across the full geographical range, using 2,406,962 SNPs (Figure 3).

**Figure 3.**
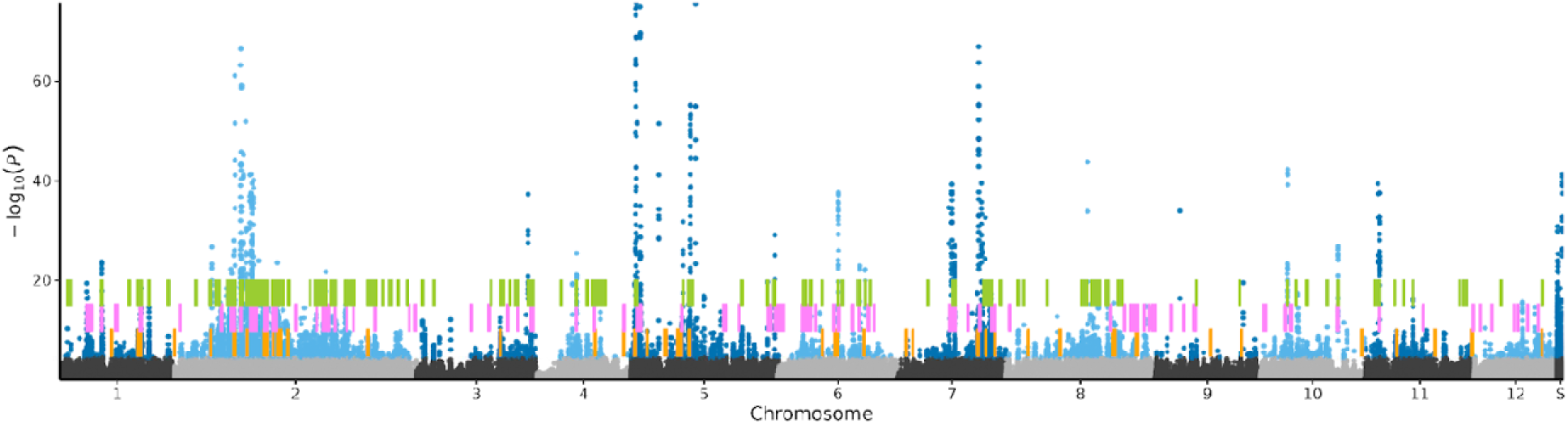
– Manhattan plot of p-values from tests for outlier loci showing excess differentiation between *Q. robur* and *Q. petraea* relative to the genome-wide background based on Dataset 3 (which excludes admixed individuals) using snmf. SNPs with a q value FDR <0.01 -. (pers. comm); Pink: Leroy et al. ^52^; Yellow - Nocchi et al. ^13^.

We found 7,930 SNPs significantly differentiated (FDR < 0.01) between the two species following sparse non-negative matrix factorisation (snmf) calculations of ancestry coefficients including all the samples in **dataset 3** (Figure 3). F_st_ values calculated in **dataset 3** for these significant SNPs ranged from 0 to 0.98. Significant SNPs were mostly found on chromosomes 2, 5, 7, 11 and regions of the unassigned scaffolds (Figure 3). Most of chromosomes 3 and 12, and particularly chromosome 9, show very little differentiation and are likely to be freely exchanging between the two species.

In **dataset 2**, we found 6,862 significantly differentiated SNPs at FDR <0.01 (Supplementary Figure 4). Of these, 6,554 SNPs were common to both **dataset 2** and **dataset 3** analyses. Of the 8,238 significant SNPs identified across both analyses, 2,005 (24%) were found inside 945 predicted genes from the *Quercus robur* reference genome^64^. A list of the significant SNP identifiers along with a table listing the predicted genes and their possible functions are available at https://github.com/biolougy/PhD. The location of highly differentiated genes found by other studies is compared with our results in Figure 3 and Supplementary Figure 4.

### Niche differentiation

We show that *Q. robur* and *Q. petraea* have clear differentiation in climate, topographic gradients, and soil preferences using PCA analyses including all environmental variables (Figure 4) and a beta regression with a subset of 13 non-collinear environmental variables (Table 2, Supplementary Figure 5). In a PCA including only climatic variables (n = 19), PC1 explained 54.9% of the variance (Figure 4a). The *Q. petraea* cluster was more loaded by precipitation variables, whereas the *Q. robur* cluster by temperature variables (Figure 4a). In the beta regression (Table 2), precipitation of the warmest quarter (Bio18, χ² = 26.94, p < 0.001) and isothermality (Bio3, χ² = 6.47, p = 0.01) were negatively associated with *Q. robur* admixture, whereas temperature seasonality (Bio4; χ² = 19.2, p < 0.001), mean temperature of the wettest quarter (Bio8; χ² = 7.86, p = 0.005), mean temperature of the coldest quarter (Bio11, χ² = 4.44, p = 0.04) and precipitation seasonality (Bio15; χ² = 12.94, p < 0.001) were positively associated with it. We found no significant association for mean temperature of the driest quarter (Bio9; χ² = 1.14, p = 0.29).

**Figure 4.**
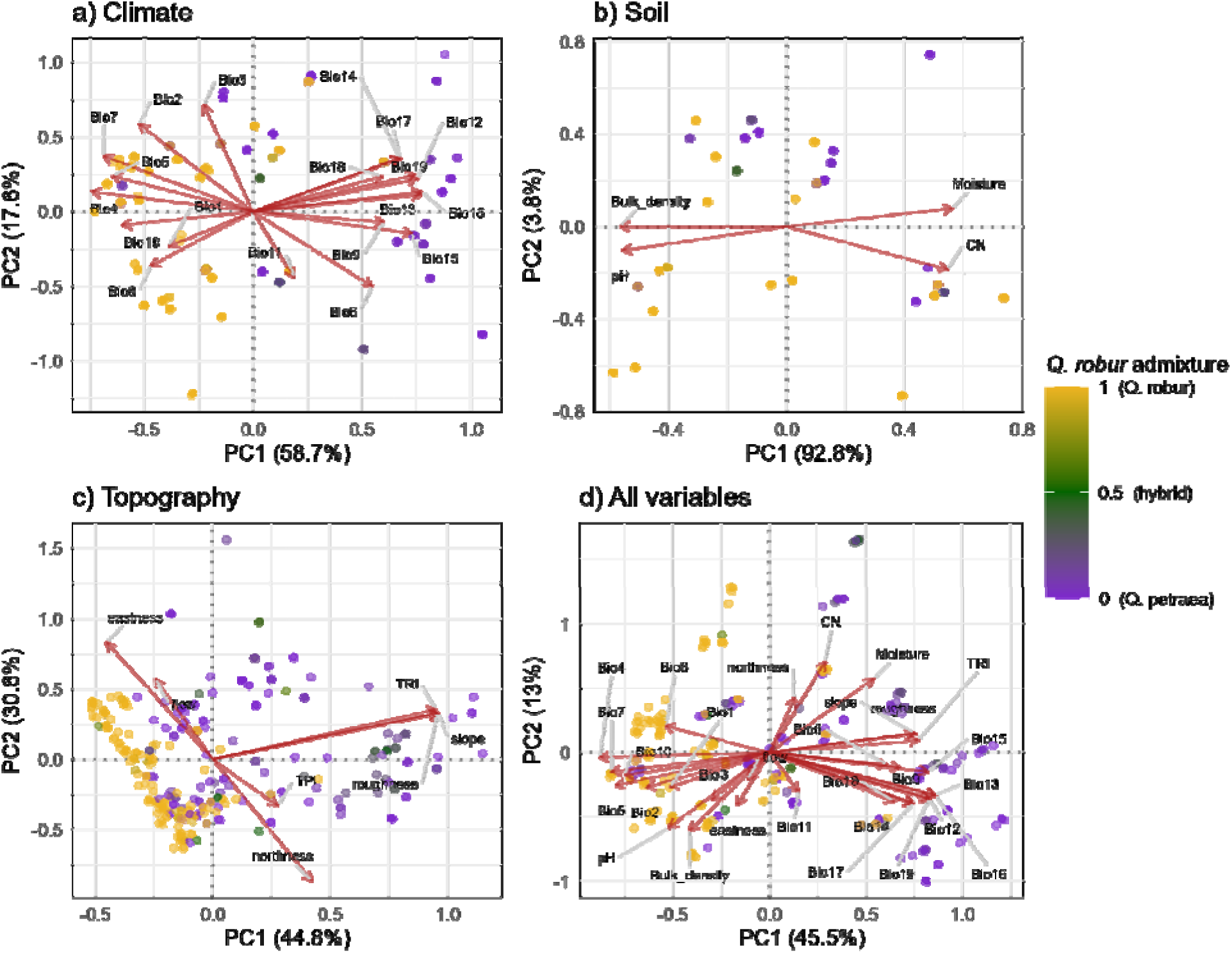
– Principal component analysis (PCA) biplots of environmental variables measured across 416 oak trees, with points coloured by *Quercus robur* admixture proportion (purple, pure *Q. petraea*; green, admixed/hybrid; gold, pure *Q. robur*). Separate PCAs were performed for a) climate variables, b) soil variables, and c) topographic variables, with a combined PCA including all variable groups shown in d). Aspect was decomposed into northness and eastness prior to analysis. Arrows indicate the direction and relative contribution of each environmental variable to the ordination space; only variables exceeding a minimum loading threshold of 0.25 are shown for clarity. Climate variables comprised the 19 standard bioclimatic variables (BIO1–BIO19): Annual Mean Temperature (BIO1), Mean Diurnal Range (BIO2), Isothermality (BIO3), Temperature Seasonality (BIO4), Maximum Temperature of the Warmest Month (BIO5), Minimum Temperature of the Coldest Month (BIO6), Temperature Annual Range (BIO7), Mean Temperature of the Wettest (BIO8), Driest (BIO9), Warmest (BIO10) and Coldest (BIO11) quarters, Annual Precipitation (BIO12), Precipitation of the Wettest (BIO13) and Driest (BIO14) months, Precipitation Seasonality (BIO15), and Precipitation of the Wettest (BIO16), Driest (BIO17), Warmest (BIO18) and Coldest (BIO19) quarters.

**Table 2.**
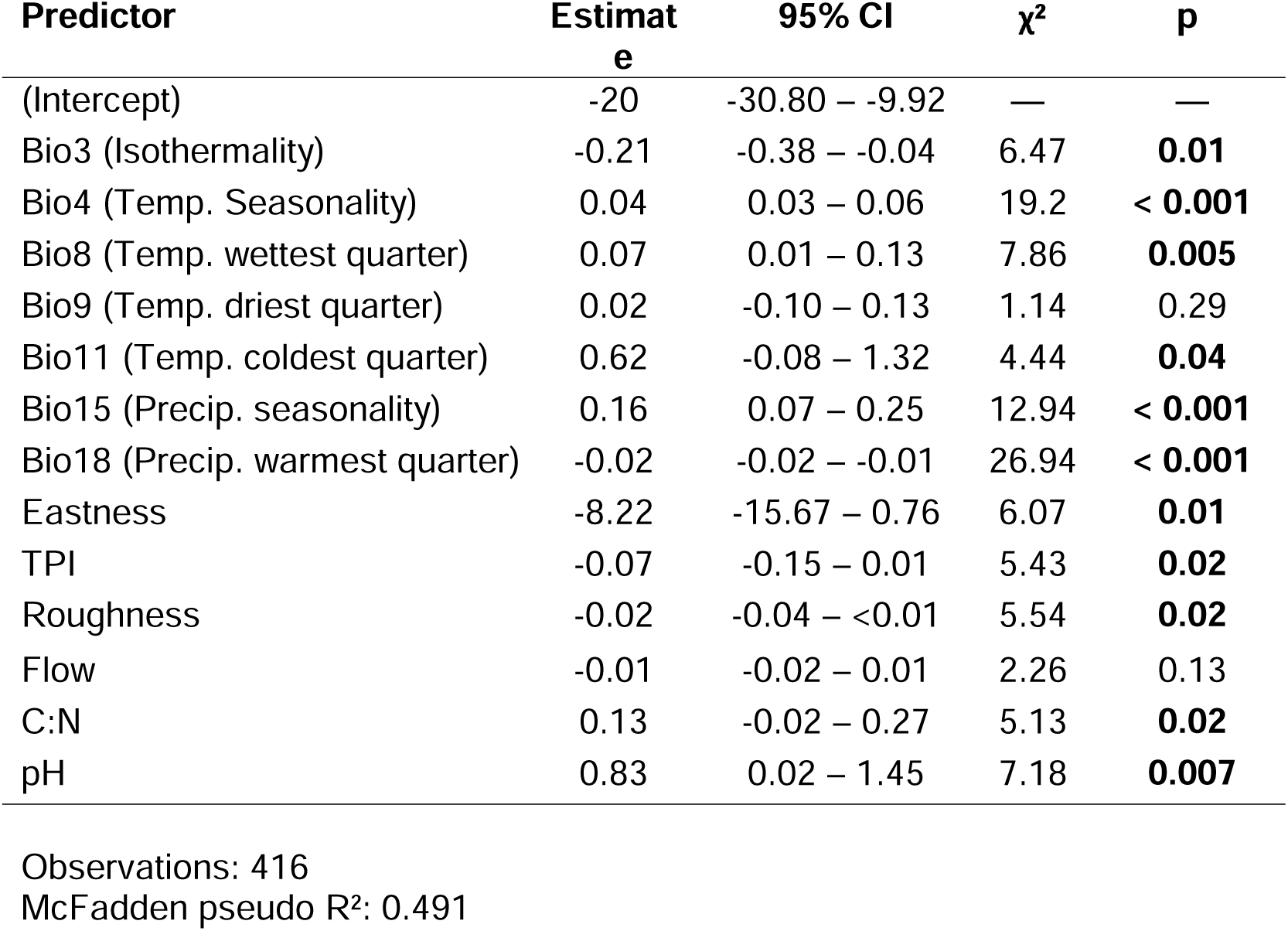
– Summary statistics for a beta regression model investigating the association between environmental predictors and *Q. robur* admixture proportion. Estimates are on the logit scale. χ² and p were estimated with likelihood ratio tests. Significant p-values are in bold. Notes: CI = confidence intervals.

In the PCA including only soil variables (n = 4), PC1 alone explained 92.8% of variance (Figure 4b), and separation between species was mostly driven by pH and soil C:N. Similarly, in the beta regression (Table 2, Supplementary Figure 5), *Q. robur* admixture was positively associated with soil pH (χ² = 7.18, p = 0.007) and C:N (χ² = 5.13, p = 0.02). For topographic variables (n = 7), greater slope, terrain roughness, and terrain ruggedness index (TRI) separated the *Q. petraea* cluster from the *Q. robur* cluster in the PC1 axis, which explained 44.5% of the variance (Figure 4c). Eastness, northness, TPI and flow drove the PC2 axis (29.9%). Similarly, eastness (χ² = 6.07, p = 0.01), TPI (χ² = 5.43, p = 0.02), and terrain roughness (χ² = 5.54, p = 0.02) were all negatively associated with *Q. robur* admixture in the beta regression (Table 2, Supplementary Figure 5), but flow accumulation was not (χ² = 2.26, p = 0.13).

The combined PCA (PC1 = 45.7%, PC2 = 12.8%, Figure 4d) confirmed the general picture that, relative to *Q. robur*, in Britain *Q. petraea* occurs at locations with rougher terrains, lower temperatures with less seasonal variation, and more acidic soils. The 13 environmental variables explained a reasonable proportion of the variance in *Q. robur* admixture (McFadden pseudo R²: 0.491, Table 2).

### Growth rates

Of the trees in the study, 322 had DBH measurements for 1990 and 2019. During this time, trees grew on average 106.7 mm (standard deviation: 53.14 mm). As a percentage of the 1990 measurement, this mean growth was 12.55% (SD = 7.53%) for all trees. Within these, percentage growth was greater for *Q. robur*, at 14.6% (N = 148; SD = 8.57%), than for *Q. petraea* at 11.1% (N = 117; SD = 6.1%) and hybrids at 10% (N = 57; SD = 5.7%) (see Figure 5a where log values are plotted). However, our linear mixed models accounting for site as random effects showed that the difference in size between species is not significant (χ² = 3.95, p = 0.139; Figure 5, Table 3).

**Figure 5.**
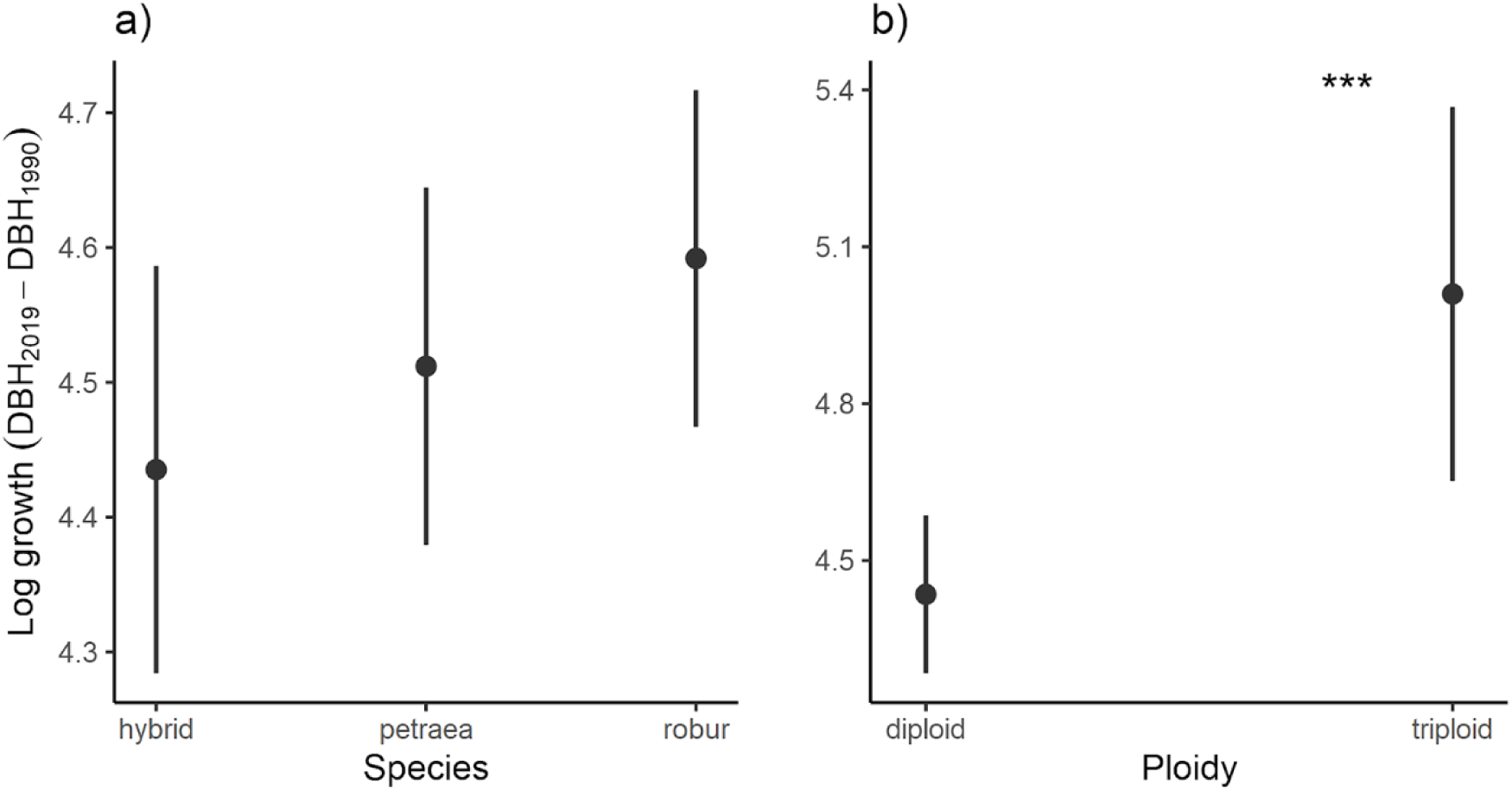
– DBH growth from 1990 to 2019 with respect to oak species and genome ploidy. Dots and error bars represent the marginal effects of the mean and the 95% confidence intervals, respectively. DBH growth was calculated as the difference in DBH between 2019 and 1990, and it was log-transformed to achieve normality of the residuals. Note: *** = p < 0.001.

**Table 3.** – Summary statistics of a linear mixed model describing the variation in DBH growth from 1990 to 2019 by oak species (i.e. Q. robur, Q. petraea, and hybrids) and individual genome ploidy (i.e. diploid, triploid). Significant p-values are in bold. Notes: CI = confidence intervals; LRT = likelihood ratio test; σ^2^ = residual variance; τ = site variance.

| Predictors | Estimate | 95% CI | $\chi^2$ | p |
| --- | --- | --- | --- | --- |
| (Intercept) | 4.07 | 3.81 – 4.32 |  |  |
| Species |  |  | 3.95 | 0.139 |
| └ *Q. petraea* | 0.08 | -0.06 – 0.22 |  |  |
| ▢ *Q. robur* | 0.16 | <0.01 – 0.31 |  |  |
| DBH<sub>1990</sub> | <0.01 | <0.01 – <0.01 | 12.67 | **<0.001** |
| Ploidy [triploid] | 0.57 | 0.25 – 0.9 | 11.83 | **<0.001** |
| **Random Effects** |  |  |  |  |
| $\sigma^2$ | 0.12 | | | |
| $\tau_{00 \text{ Site}}$ | 0.11 | | | |
| ICC | 0.49 |  |  |  |
| N<sub>Site</sub> | 47 |  |  |  |

Observations: 322
Marginal R<sup>2</sup> / Conditional R<sup>2</sup>: 0.08 / 0.531

Accordingly, the fixed effects of ploidy and species only explained ∼8% of the variance in growth whereas fixed and random effects combined explained ∼53% of the variance (Table 3). However, we found that triploid individuals grew on average more (16.6%, SD = 5.68%) than diploid individuals (12.5%, SD = 7.53%), even when site variation was accounted for (χ² = 11.83, p < 0.001; Figure 5, Table 3). This relationship remained significant even if we restricted the analysis to sites where both ploidy types co-occurred (χ² = 6.6, p = 0.010). One triploid was the third fastest-growing tree of the 322, with growth of 296 mm in 29 years.

To identify the most important environmental variables contributing to tree growth, we removed the random effect of site identity and ran a linear model using the same uncorrelated climatic, soil, and terrain variables in the previous analysis (Table 4). We found that, in addition to ploidy, growth was positively associated with isothermality (F_1,301_ = 5.62, p = 0.018; Figure 6a) and mean temperature of the coldest quarter (F_1,301_ = 4.21, p = 0.041; Figure 6b), and negatively associated with terrain roughness (F_1,301_ = 9.38, p = 0.002; Figure 6c). The full model explained ∼21-26% of the variance in growth (Table 4). The lower variance explained by this model compared to the linear mixed model suggests other unmodelled site-specific environmental variables may play an important role in oak growth.

**Figure 6.**
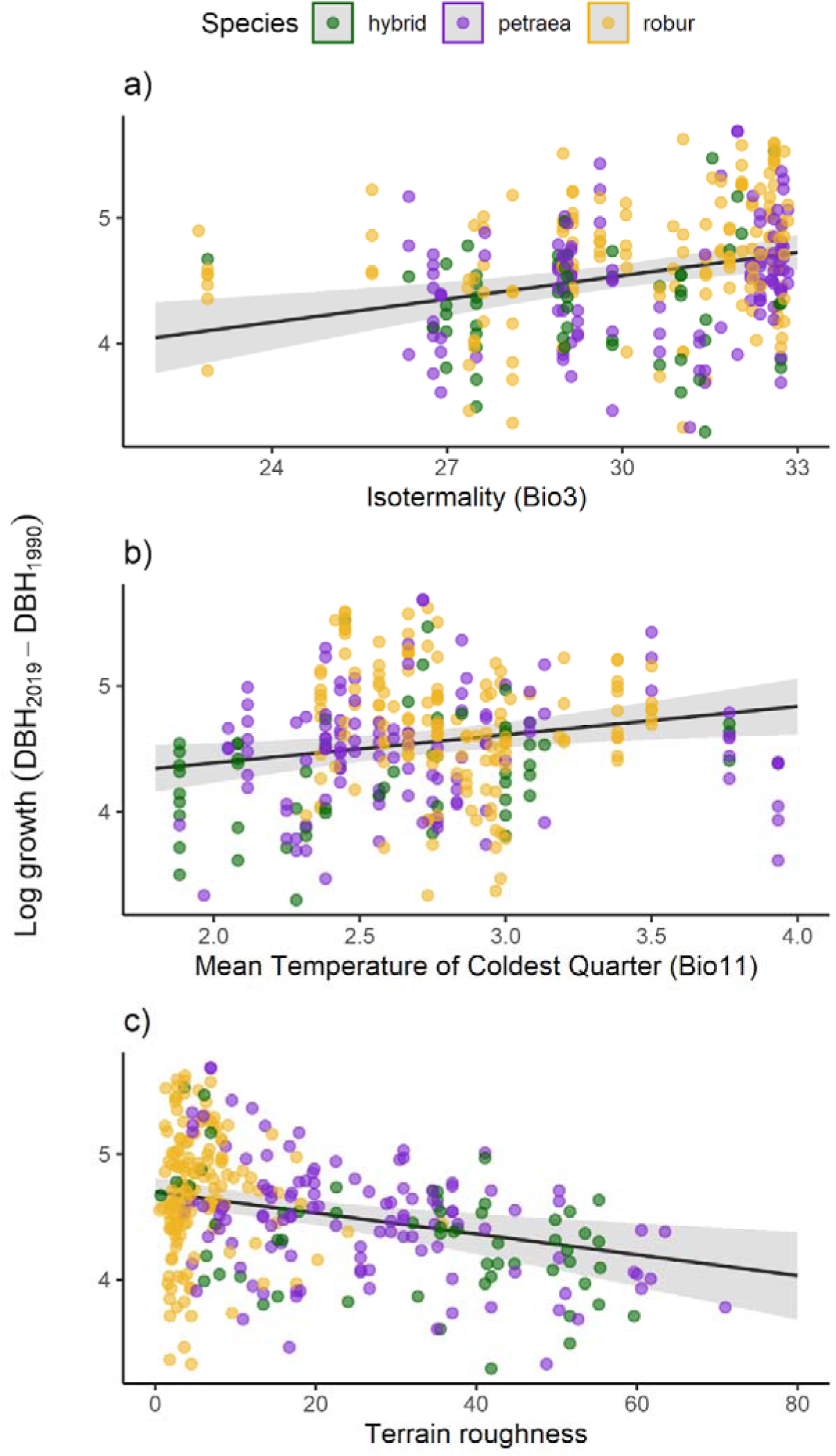
– DBH growth from 1990 to 2019 with respect to site environmental conditions. Black lines and shades represent the marginal effects and the 95% confidence intervals, respectively. Dots represent the raw datapoints, which are coloured by species for visualisation purposes only (gold = *Q. robur*; purple = *Q. petraea*; green = hybrids). Growth was calculated as the difference in DBH between 2019 and 1990, and it was log-transformed to achieve normality of model residuals. Only significant associations are shown (see Table 4).

**Table 4.** – Summary statistics of a linear model describing the variation in DBH growth from 1990 to 2019 by environmental, terrain, and soil characteristics. Oak species (i.e. *Q. robur*, *Q. petraea*, and hybrids) and individual genome ploidy (i.e. diploid, triploid) were entered as covariates. Significant p-values are in bold. Notes: CI = confidence intervals.

| Predictors | Estimate | 95% CI | F-statistic | p |
| --- | --- | --- | --- | --- |
| (Intercept) | 2.73 | -0.06 – 5.52 |  |  |
| Species |  |  | 0.093 | 0.911 |
| └─ *Q. petraea* | 0.02 | -0.13 – 0.18 |  |  |
| └─ *Q. robur* | 0.04 | -0.14 – 0.22 |  |  |
| DBH<sub>1990</sub> | <0.01 | <0.01 – <0.01 | 2.663 | 0.104 |
| Ploidy (triploid) | 0.61 | 0.22 – 0.99 | 9.544 | **0.002** |
| Bio3 (Isothermality) | 0.06 | 0.03 – 0.10 | 13.062 | **0** |
| Bio4 (Temp. Seasonality) | <0.01 | -0.01 – <0.01 | 0.941 | 0.333 |
| Bio8 (Temp. wettest quarter) | <0.01 | -0.01 – 0.02 | 0.091 | 0.763 |
| Bio9 (Temp. driest quarter) | 0.01 | -0.02 – 0.03 | 0.146 | 0.703 |
| Bio11 (Temp. coldest quarter) | 0.22 | 0.06 – 0.39 | 7.13 | **0.008** |
| Bio15 (Precip. seasonality) | -0.02 | -0.04 – 0.01 | 1.744 | 0.188 |
| Bio18 (Precip. warmest quarter) | <0.01 | <0.01 – <0.01 | 0.021 | 0.886 |
| Aspect | 0.02 | -0.01 – 0.05 | 1.912 | 0.168 |
| Topographic Position Index | 0.01 | -0.01 – 0.03 | 0.857 | 0.355 |
| Roughness | -0.01 | -0.01 – <0.01 | 10.367 | **0.001** |
| Flow accumulation | <0.01 | <-0.01 – < 0.01 | 0.404 | 0.526 |
| C:N ratio | 0.02 | -0.01 – 0.06 | 2.055 | 0.153 |
| Soil pH | 0.05 | -0.07 – 0.18 | 0.697 | 0.404 |

Observations: 322
R<sup>2</sup> / R<sup>2</sup> adjusted: 0.258 / 0.216

**Table 5.**
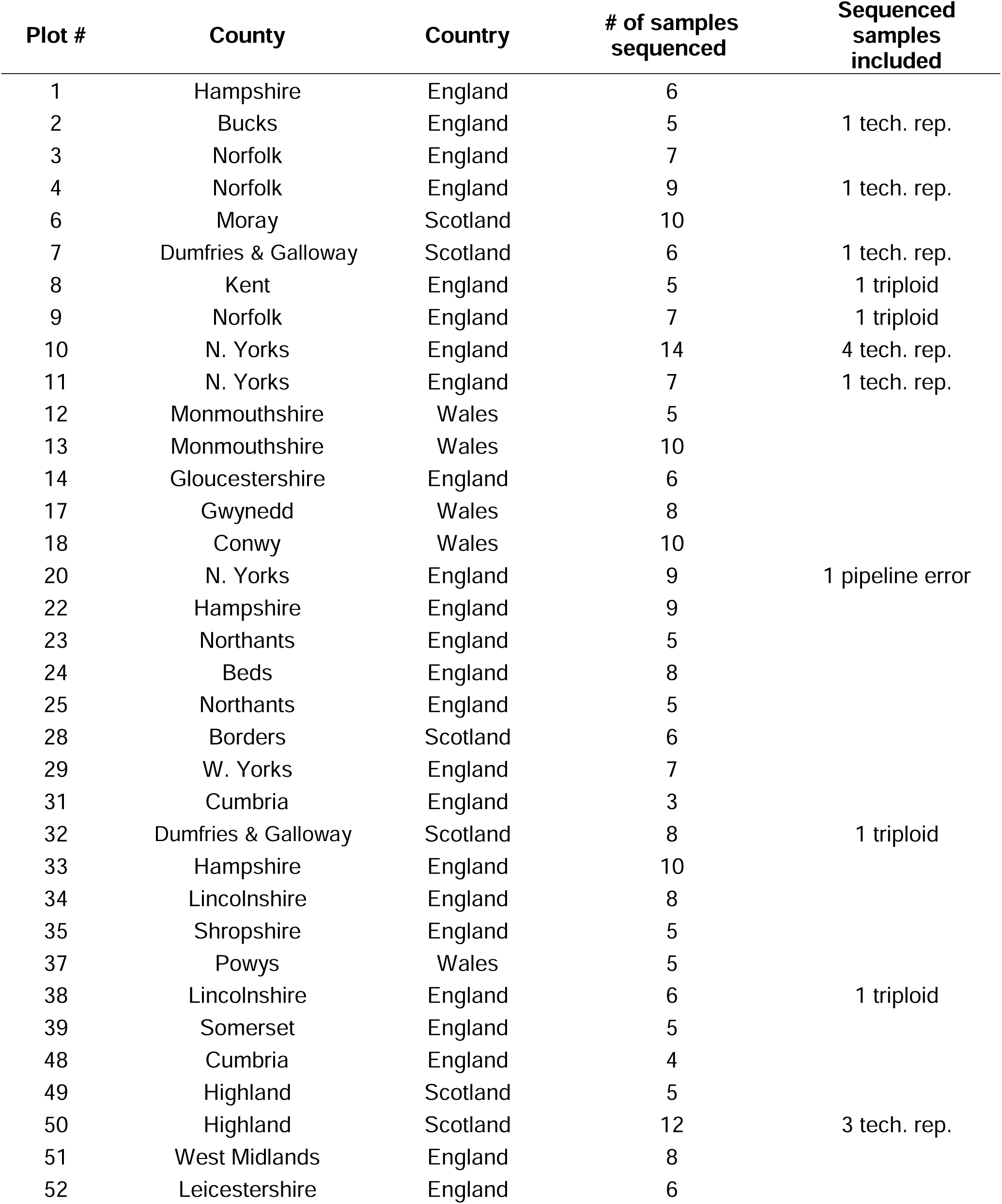
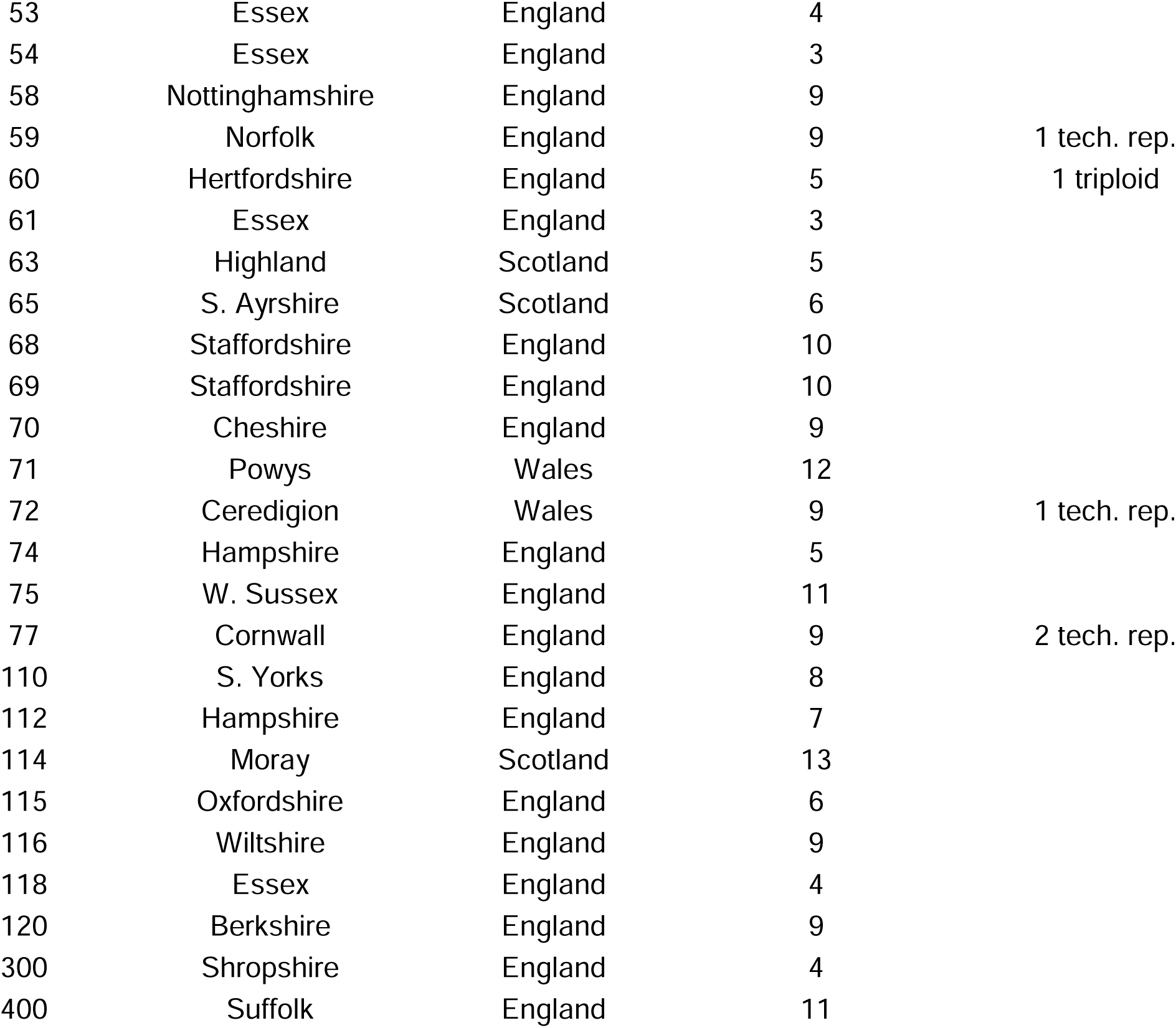
– Country and county locations, woodland type and number of samples from each woodland (total = 418 samples).

| Plot # | County | Country | # of samples sequenced | Sequenced samples included |
| --- | --- | --- | --- | --- |
| 1 | Hampshire | England | 6 |  |
| 2 | Bucks | England | 5 | 1 tech. rep. |
| 3 | Norfolk | England | 7 |  |
| 4 | Norfolk | England | 9 | 1 tech. rep. |
| 6 | Moray | Scotland | 10 |  |
| 7 | Dumfries & Galloway | Scotland | 6 | 1 tech. rep. |
| 8 | Kent | England | 5 | 1 triploid |
| 9 | Norfolk | England | 7 | 1 triploid |
| 10 | N. Yorks | England | 14 | 4 tech. rep. |
| 11 | N. Yorks | England | 7 | 1 tech. rep. |
| 12 | Monmouthshire | Wales | 5 |  |
| 13 | Monmouthshire | Wales | 10 |  |
| 14 | Gloucestershire | England | 6 |  |
| 17 | Gwynedd | Wales | 8 |  |
| 18 | Conwy | Wales | 10 |  |
| 20 | N. Yorks | England | 9 | 1 pipeline error |
| 22 | Hampshire | England | 9 |  |
| 23 | Northants | England | 5 |  |
| 24 | Beds | England | 8 |  |
| 25 | Northants | England | 5 |  |
| 28 | Borders | Scotland | 6 |  |
| 29 | W. Yorks | England | 7 |  |
| 31 | Cumbria | England | 3 |  |
| 32 | Dumfries & Galloway | Scotland | 8 | 1 triploid |
| 33 | Hampshire | England | 10 |  |
| 34 | Lincolnshire | England | 8 |  |
| 35 | Shropshire | England | 5 |  |
| 37 | Powys | Wales | 5 |  |
| 38 | Lincolnshire | England | 6 | 1 triploid |
| 39 | Somerset | England | 5 |  |
| 48 | Cumbria | England | 4 |  |
| 49 | Highland | Scotland | 5 |  |
| 50 | Highland | Scotland | 12 | 3 tech. rep. |
| 51 | West Midlands | England | 8 |  |
| 52 | Leicestershire | England | 6 |  |
| 53 | Essex | England | 4 |  |
| 54 | Essex | England | 3 |  |
| 58 | Nottinghamshire | England | 9 |  |
| 59 | Norfolk | England | 9 | 1 tech. rep. |
| 60 | Hertfordshire | England | 5 | 1 triploid |
| 61 | Essex | England | 3 |  |
| 63 | Highland | Scotland | 5 |  |
| 65 | S. Ayrshire | Scotland | 6 |  |
| 68 | Staffordshire | England | 10 |  |
| 69 | Staffordshire | England | 10 |  |
| 70 | Cheshire | England | 9 |  |
| 71 | Powys | Wales | 12 |  |
| 72 | Ceredigion | Wales | 9 | 1 tech. rep. |
| 74 | Hampshire | England | 5 |  |
| 75 | W. Sussex | England | 11 |  |
| 77 | Cornwall | England | 9 | 2 tech. rep. |
| 110 | S. Yorks | England | 8 |  |
| 112 | Hampshire | England | 7 |  |
| 114 | Moray | Scotland | 13 |  |
| 115 | Oxfordshire | England | 6 |  |
| 116 | Wiltshire | England | 9 |  |
| 118 | Essex | England | 4 |  |
| 120 | Berkshire | England | 9 |  |
| 300 | Shropshire | England | 4 |  |
| 400 | Suffolk | England | 11 |  |

## Discussion

We resequenced the whole genomes of 423 oak individuals sampled from a long-term survey spanning 60 sites across the United Kingdom to comprehensively characterise population structure in British oaks. We show that the two native species *Quercus robur* and *Quercus petraea*: (1) have clear geographic differentiation and niche preferences, (2) are freely admixing particularly in Scotland, (3) are differentiated in many regions of the genome, and (4) each contain a few putative triploid individuals with high growth rates. Our findings suggest that the continuum between the two British oak species is maintained by the opposing forces of hybridisation and niche differentiation. They challenge some aspects of the “resurrection” model^15^ for the colonisation of Britain

This study contributes to our understanding of the differentiation of *Q. robur* and *Q. petraea* in a corner of their native range where they have influenced the development of evolutionary biology. We found a clear pattern that *Q. robur* is more common in the south and east, and *Q. petraea* more common in the north and west. This is broadly in agreement with old reports of the distribution of each species in Britain^5–7,11,30–32^,. Unlike previous Britain-wide studies we were able to distinguish between the two species on the basis of whole genome data, rigorously measure the levels of admixture within hybrids, and conduct correlations with quantitative environmental data. We found a clear relationship between the proportion of the genome that is *Q. robur*, and a more continental climate (i.e. lower isothermality and higher temperature seasonality), lower soil acidity and in less rough and complex terrain. This confirms conclusions from many observational studies of the apparent habitat preferences of the two morphological species, both in Britain and on the continent^5,30,31,33–35,41,42^ and also fits with physiological studies^39,42–46^.

Despite their clear environmental preferences, we find considerable evidence for ongoing hybridisation between *Q. robur* and *Q. petraea* throughout Britain. We found several F_1_ hybrids and a continuum of levels of introgression attributable to back crossing, mainly with *Q. petraea* but occasionally with *Q. robur*. These findings are consistent with previous studies from other areas of northern Europe, which also found widespread hybridization^12,15,49,50,52,65–67^. It also confirms earlier suggestions of extensive introgression in Scotland from nineteenth century naturalists, and twentieth studies demonstrating extensive variation in leaf morphology^3,4,6^. A recent study in Ireland found even higher levels of introgression between the two species, albeit based on only 441 loci^68^.

One interpretation of latitudinal and environmental clines of gene flow between *Q. robur* and *Q. petraea* is that introgression provides alleles that are adaptive to environments outside the niche of each species^15,52^. At two Scottish sites (numbers 28 and 63), all samples were F_1_ hybrids or back-crossed *Q. petraea*, apart from a single back-cross with *Q. robur*. Both sites are on steep slopes, so it is possible that introgression from *Q. petraea* may allow *Q. robur* to adapt to rougher terrain than is normally found^47^. Carleial et al.^69^ have shown tentative evidence that introgression from *Q. petraea* could help *Q. robur* develop some resistance to powdery mildew, but as this disease is a recent introduction, selection from it cannot have contributed much to genetic patterns in adult trees. Introgression adaptive to novel environments could have occurred early during postglacial colonisation, involving alleles that are now fixed in both species and therefore difficult to detect within Britain alone. Broader European-scale analyses may help resolve these possibilities.

An alternative, neutral, hypothesis would be that the pattern we see of greater introgression in the north is due to a greater propensity for hybridisation in colder environments. A local account from the Botanical Society of Britain and Ireland suggests increased flowering overlap between *Q. petraea and Q. robur* at higher elevations in County Fermanagh (https://bsbi.org/in-your-area/local-botany/co-fermanagh/fermanagh-species-accounts/quercus-petraea-liebl). In birch species, it has long been suggested that low temperatures in the north may facilitate hybridization^70^, and experiments on *Arabidopsis* hybrids show that lower temperatures during seed development increase the survival of hybrid F_1_ seeds^71^. Thus, it is possible that oaks could have a greater propensity to be more able to hybridise at high latitudes and altitudes.

We also found several regions of the genome with little evidence of introgression. These comprised 8,238 statistically significant SNPs and 945 predicted genes, 21 of which were in common with the 84 genes found by Nocchi et al.^13^ in England and 34 were in common with the 127 genes identified by Leroy et al.^49^. A central European study, using a similar approach to ours, has recently found 249 genes differentiating the two species, 153 of which are also found amongst the 945 identified here (Lazic, pers. com). The degree of overlap between British and continental studies suggests a degree of consistency in the species boundary across Europe, and that the species differences between *Q. robur* and *Q. petraea* are due to many locations in the genome, not a few. In general, the genus *Quercus* is known to exhibit widespread ongoing hybridisation, without leading to species diversification or affecting species integrity^19^.

Our results are difficult to reconcile with the “resurrection” model, which proposes that *Q. robur* acted as the primary postglacial coloniser of Europe, with *Q. petraea* subsequently expanding through pollen swamping of pre-existing *Q. robur* populations^15^. As oaks colonised Britain following the last glacial maximum primarily from southern and eastern refugia^7,25,26^, the resurrection model would predict greater representation of *Q. petraea* in southern and eastern Britain and *Q. robur* in northern and western regions. The model also posits that a *Q. petraea* phenotype can emerge rapidly within a predominantly *Q. robur* genomic background through introgression at only a small number of loci. However, we found the opposite geographical distribution pattern and many differentiated loci. Although introgression in our dataset is asymmetrical, the magnitude of this bias does not appear sufficient to support extensive historical pollen swamping. Instead, it may reflect modest but persistent asymmetry in hybridisation potential between the species. We also detected limited backcrossing into *Q. robur*, indicating bidirectional gene flow. The high prevalence of hybrids in Scotland may partly reflect historical planting of *Q. robur* into landscapes previously dominated by *Q. petraea*. At the same time, the geographic concentration of introgression in regions where both species co-occur argues against ancestral shared polymorphism as the primary explanation for allele sharing^72^.

Reconciling our results with the resurrection model would require *Q. robur* to have initially colonised the entire island, followed by replacement by *Q. petraea* only in northern and western regions. Such a scenario would imply strong regional selection against *Q. petraea* in southern and eastern Britain, while also requiring long-distance dispersal across areas where the species was apparently disfavoured. An alternative explanation is that pollen swamping played a more limited role in determining present-day species distributions, and that *Q. petraea* colonised northern and western Britain directly through acorn dispersal, followed by environmental selection favouring its persistence in these regions.

The chloroplast haplotypes provide an additional perspective on postglacial recolonisation because chloroplast DNA in oaks is maternally inherited and therefore reflect acorn-mediated dispersal rather than pollen flow. Most of our samples were in nodes that correspond to restriction enzyme haplotypes 10, 11, and 12 from chloroplast lineage B. This lineage is common from Spain to Denmark, and has been previously identified as the most common chloroplast lineage in Britain^11,21^. It is thought to have originated from a Late Pleistocene glacial refugium in the Iberian peninsula^63^. Lineage C, which we putatively find in two individuals from Hazelborough wood, is widespread in central western Europe and is thought to have an Italian glacial refugium^21^. Lineage A, which we find in nine trees in Gloucester and Chester, is widespread in France and central Europe^21^. The glacial refugium for this haplotype could be Spain, Italy or the Balkans, with Petit et al favouring the latter hypothesis^63^. All the haplotype lineages that we found have previously been found in Britain. As with previous studies we did not find them to be differentiated among species. All detected haplotypes were consistent with natural post-glacial recolonisation from Europe.

Our study is unusual for a population genomic study in that we have access to long-term growth rate data for many samples. We found *Q. robur* to be growing faster than both *Q. petraea* and hybrids, but this was mainly attributable to the environmental conditions in which they are growing, particularly isothermality, temperature of the coldest quarter and the roughness of the terrain. Early results from oak provenance trials in Britain^73^ did not find clear trends in the growth rate of young *Q. robur* versus *Q. petraea*, but did note that the best *Q. robur* provenances performed less well in the most northerly and western trial sites. In high forests of central and northwestern France, *Q. petraea* grows significantly faster than *Q. robur*^74^, but early results from provenance trials in the Czech Republic suggest that *Q. robur* may grow faster in its typical site conditions^75^. An experiment on one year old seedlings from Germany showed faster growth by *Q. robur*^76^. In the past, planting of *Q. robur* has often been recommended or practised in Scotland^5,34,36,37^, but this may have been based on observed growth rates of *Q. robur* in more favourable environments. Our results allow careful selection of sites suitable for each species and their hybrids.

Interestingly, we found evidence for five triploid individuals amongst our samples in both *Q. robur* and *Q. petraea* dominated woodlands, and that these have significantly higher growth rates than diploids even when environmental variables are accounted for. These triploids could be cloned for high carbon capture plantations. Our genomic analyses suggest that two triploids are *Q. robur*, two are *Q. petraea* and one is a back-crossed admixed individual that is predominantly *Q. petraea*. This fits with previous discoveries of triploids in both species^77–79^ Occasional triploid individuals have been reported from several European populations^77–79^. These individuals were associated with unusual morphology, reduced fertility, or enlarged stomata.

## Conclusion

When Darwin referred to the lack of differentiation between British oaks, he was making a case that varieties and species form a continuum that arises via descent with modification. Although he did not claim that British oaks would continue to differentiate until they became distinct and unambiguous species, he clearly viewed this as a possibility, and one that must have happened innumerable times in other cases even if specific circumstances prevented it in this instance. Here, we show in genomic detail how a complex interaction of niche differentiation and extensive gene flow maintain a dynamic yet apparently stable complex in which *Q. robur* and *Q. petraea* coexist as distinct forms, yet continue to intergrade. The presence of two species with different environmental preferences and the ability to share and exchange adaptive variation has no doubt contributed greatly to the success of oak as the most common broadleaf species in Britain.

## Acknowledgements

The Defra Future Proofing Plant Health programme provided funding for sequencing all of the collected samples. Samples were collected and phenotyped by staff of the FR TSU, including: Liz Richardson, Steve Whall, Matthew Anstey, Duncan Williams. Woodland Heritage funded Nathan Brown’s time. Louise Gathercole’s PhD was supported by a studentship funded by QMUL and an Action Oak grant to RBG Kew.

## Author Contributions

Nathan Brown provided sample meta-data and coordinated collecting with the FR TSU. Gabriele Nocchi helped process incoming samples at Kew and contributed to the chloroplast analyses. Sandra Denman suggested the study and helped in its conceptualisation and initiation. Louise Gathecole analysed the data and wrote the first draft of the MS. Richard Buggs obtained funding and oversaw the project. Richard Buggs and Richard Nichols co-supervised Louise Gathercole’s PhD project. Rômulo Carleial and Ernest Wu contributed analyses. Louise Gathercole, Rômulo Carleial, Ernest Wu and Richard Buggs wrote the final draft of the MS.

## Data Availability

Novel read data generated in this study are available at: https://www.ebi.ac.uk/ena/browser/view/PRJEB43909

Results from the Forest Condition Survey are available at: https://www.data.gov.uk/dataset/cccef1ac-fcd2-456a-89eb-49f7206e9ce1/forest-condition-survey-1987-2006

## Materials and Methods

### Sample collection

The vast majority of the sampled trees were in plots (Table 5) that originally formed part of the Forest Condition Survey by Forest Research, which ran between 1987 and 2007 as part of the European ICP initiative (The International Co-operative Programme on Assessment and Monitoring of Air Pollution Effects on Forests) documenting the health of five tree species ^80^. In 1990, the survey assessed 7,644 trees over 319 sites, 73 of which were oak plots, and site selection was designed to achieve an even coverage in the main areas of Britain where oak is found ^80^. The oak plots were first surveyed in 1987 ^62^. In each of 12 regions of Britain, three plots with 24 trees were sought: one for each of the age classes 80-110, 111-140 and 141-180 years ^62^. Each selected stand had to have no edge trees, no windthrow, and all 24 crowns had to be visible from the ground. Such stands could not be found in some parts of Scotland due to the rarity of stands of the appropriate age ^62^. Surveys were initially set up to monitor forest condition in response to concerns about the effects of acid rain on woodlands: this included measures of diameter at breast height (DBH) and crown condition. The value of such surveys for monitoring the responses of trees to environmental stressors such as drought, pests and pathogens, and to climate changes was recognised and monitoring continued in Britain until 2007 when funding ceased ^80^. Data from the study was recently made available here: https://www.data.gov.uk/dataset/cccef1ac-fcd2-456a-89eb-49f7206e9ce1/forest-condition-survey-1987-2006 (URL accessed 17 December 2025). Following the Action Oak Knowledge Review (see https://www.actionoak.org/s/Action-Oak-Knowledge-Review-12-06-2019.pdf), commissioned in 2019, funding was provided by the Forestry Commission to revisit documented oaks at some of the plots and this was taken as an opportunity to collect leaf samples in case of future opportunities to carry out sequencing across the range of oaks in Britain. In the summer of 2019 the Forest Research Technical Services Unit located and assessed 85 plots using the ICP Forests methodology (see https://www.icp-forests.net/monitoring-and-research/icp-forests-manual). The survey took place between June and October. During the survey, leaf samples were collected from 761 trees at 66 of the sites. Samples were placed in sealed bags with silica gel and sent to the Royal Botanic Gardens, Kew where they were dried in the bags at room temperature. For this study, sample sites were selected from the survey to cover a broad geographical area and with the aim of having at least four samples available at eachthe site. The DBH of each tree was measured. Woodland type was assessed using the site woodland management plans where available or the ‘Magic map’ (available from https://magic.defra.gov.uk/MagicMap.aspx ©Crown copyright and database rights). Details of county and country and the number of samples for each of the sites can be seen in Table 5. Two of the sites (300 and 400) were not part of the original Forest Condition Survey, but are part of a separate monitoring project looking at acute oak decline ^69,81^.

### DNA extraction and sequencing

We selected 10mg per each of 462 dried leaf samples which were sent to Novogene for DNA extraction and resequencing. DNA was extracted using the EchoLUTION Plant DNA 96 Core Kit (BioEcho Life Sciences, Cologne, Germany) protocol with the following modifications: A proprietary additive was added during the lysis step to remove DNA blocking inhibitors. 10µl of 3M Sodium acetate pH 5.0 was added to the mix prior to loading it to the column to prevent chlorophyll from binding. Eluates were re-purified using the Qiagen QIAquick PCR purification kit (Qiagen, Hilden, Germany) with modifications: Buffer PB and 200µk of ethanol were added to the sample. Two washing steps were carried out to maximise purity. The first with 800µl of PE for 1 minute at 8000xG and the second with 400µl PE for 3 minutes at 20000xG. The ethanol in the washing buffer removed the green chlorophyll. Elution was carried out as per the kit protocol.

Genomic DNA was randomly fragmented to 350bp fragments which were end-polished, A-tailed, ligated with sequencing adapters and PCR enriched with primers of P5 and P7 oligos. The PCR libraries were purified (AMPure XP system, Beckman Coulter, Brea, California, USA) and quality control tested. Libraries were sequenced with 150-bp paired-end Illumina (San Diego, California, USA) NovaSeq 6000 technology to 20x coverage. Adapter sequences and very low quality reads were removed with fastp ^82^. In total, 425 samples plus 15 technical replicates were successfully sequenced.

### Genotyping

The reads were scanned for an average Phred base quality score of at least 20 in windows of four bases using Trimmomatic v.0.36 ^83^, and trimmed to remove the low scoring bases. Reads with fewer than 70 bases after trimming were removed. The remaining, high quality, reads were aligned to the haploid *Q. robur* reference genome ^64^ using BWA-MEM v.0.7.17 ^84^ with default settings. Putative polymerase chain reaction (PCR) duplicates were removed with Samtools v.1.9 ^85^. The Genome Analysis Toolkit (GATK) v.4.0.8.1 was used for variant calling ^86^. One sample failed to run, leaving 424 samples and 15 technical replicates. The HaplotypeCaller algorithm was used to call variants individually for each sample and create genome variant call format files ^87^. The variants were then called from each file to create a joint genotyping analysis with the GenotypeGVCFs algorithm. Single nucleotide polymorphisms (SNPs) were selected from this analysis and filtered with GATK VariantFiltration and SelectVariants. Loci were excluded if they had scores of quality-by-depth less than two, strand-odds-ratio (a measure of strand bias) greater than four, root-mean-square-mapping-quality less than 40, mapping-quality-rank sum less than -2.5 or read-position-rank-sum-test less than - 2. Filtering parameters were selected following the guidance given by the developers of GATK and based on the distribution of values within the dataset.

SNPs located in the sections of the genome identified as transposable elements were excluded using bedtools v.2.28.0 ^88^. We used BCFtools v.1.13 ^85^ to annotate the SNPs, select only biallelic SNPs and filter out SNPs with mapping quality less than 30 and individual depth less than eight or greater than 36. Parameters were selected following an analysis of the distribution of sequencing depth in the dataset.

Samples were assessed for normal allelic balance in R v.4.1.1 ^89^. Inbreeding coefficients (F) were calculated in vcftools v.0.1.16 ^90^ using a randomly selected subset of 24,297 loci. F is the reduction in observed heterozygosity relative to that expected under Hardy–Weinberg equilibrium i.e. 1 - H_O_/H_E_. Low values of F indicate excess levels of heterozygosity and could be indicative of polyploidy or sequencing errors, whereas high F values could indicate inbred individuals or individuals with high missing data. Allele balance is the proportion of sequenced reads supporting the reference allele compared with the proportion supporting the alternate allele at each locus.

Variance in allele balance of bi-allelic SNPs can be used to detect sequencing errors and is a useful quality control tool for genomic analysis. Allele balance is also a reliable metric to identify departures from diploidy ^91^. In diploids, most heterozygous loci will have close to a 0.5/0.5 balance of alleles. This indicates two haplotypes. Individuals where many loci have an allele balance 0.33/0.67 suggests triploidy ^91,92^. Sampled individuals with both negative F values and a majority of loci with allele balance of 0.33/0.67 were considered to be putative triploids, as this would indicate three versions of a locus. A fastSTRUCTURE analysis ^93^, as described in the next section, was run on all sequenced samples, which enabled us to identify the triploid species. Putative triploids were then removed from the data for all further filtering and analysis steps.

After removing triploids, heterozygous SNPs with allele balance less than 0.333 were filtered out with FilterVcf/ SelectVariants in GATK v.4.1.1. Finally, Plink v.1.9 ^94^ was used to filter SNPs for missingness of more than ten percent, and minor allele frequency of 0.002 (equivalent to two individuals in the data having the minor allele to avoid removing rare alleles). The SNP set was pruned for linkage disequilibrium (LD) with the Plink indep-pairphase function, a window size of 50 SNP markers, steps of 5 and filtering out SNPs with r^2^ > 0.4. This filtered set was then used to create a principal components analysis (PCA) summarising variation in the SNP data. We refer to this SNP set as **dataset 1**.

### Population structure, species assignment and identification of hybrids

The Forest Condition Survey meta-data does not distinguish between the two species, but records them both as ‘native oak’. To assign samples to species and to check for underlying population structure, we used fastSTRUCTURE v.1.0 ^93^ on **dataset 1**, run with a simple prior and fivefold cross-validation and the number of ancestral populations (K) from one to five. The Python function ChooseK was run on the fastSTRUCTURE outputs. This function outputs two values for K: the log-marginal likelihood lower bound of the data, which aims to identify strong structure and a second that identifies weak structure by reporting model components that have a cumulative ancestry contribution of at least 99% ^93^. The admixture coefficient (q) calculated in fastSTRUCTURE for K=2 was used to allocate a species to each sampled individual in three groups: Species 1 (q >= 0.9), admixed (q > 0.1 and q < 0.9), species 2 (q =< 0.1) following the simulations of Neophytou ^95^. Species 1 was allocated to *Q. robur* and species 2 to *Q. petraea* on the basis of the leaf morphologies of a subset of samples following guidelines described in Lemaire ^96^. There were also three individuals of known species from previous sampling whose leaf morphologies were used to inform species allocation ^13,97^. Although Neophytou ^95^ found that there was some overlap between F1 hybrids and backcrossed individuals in the value of q, we also split the hybrids into groups with q coefficient values between 0.375 and 0.625 considered to be possible F1 hybrids, 0.1 to 0.375 and 0.625 to 0.9 backcrosses. We plotted the PCA eigenvectors from the Plink output using the R package ggplot2 ^98^, coloured by the species assignment from fastSTRUCTURE as a visual assessment of population structure. To check for closely related individuals in the data, and to confirm technical replicates, we ran a relatedness analysis using vcftools v.0.1.1 -relatedness function ^90^.

### Chloroplast haplotype analysis

We assembled the chloroplast of all *Q. robur*, *Q. petraea*, and hybrid samples using NOVOPlasty 4.3.5 ^99^. The assembly was performed de novo with a kmer length of 33bp, read length of 151bp, and insert size of 300bp. We used the *Q. robur* rbcL sequence (Genbank accession # AB125025.1) as the seed sequence. The fully assembled sequences were linearized and shifted to the same origin, which was the seed sequence, using the Perl package fasta-tools (https://github.com/b-brankovics/fasta_tools) and aligned using MAFFT v7 ^100^. We then selected the most reliable regions of the alignment for haplotype network construction by removing a poorly aligned region of inverted repeats of about 22,500bp, and trimAl ^101^ was used to remove spurious sequences. We then exported the alignment in Phylip format and used PopArt v1.7 ^102^ to construct the haplotype network using the median-joining network method, with the species composition as a trait.

European oak chloroplast lineages were previously characterized by Petit et al. ^63^ using PCR restriction fragment length polymorphism (PCR-RFLP). We replicated, *in silico*, the PCR-RFLP assays for three chloroplast DNA fragment–restriction enzyme combinations described by Petit et al. (2002) for representative sequences from each of the six clusters identified in the median-joining network. Cluster identities were then matched to the established oak chloroplast haplotypes defined by Petit et al. (2002), following an approach implemented by Nocchi et al. (2022).

The chloroplast fragments and restriction enzymes analyzed were *trnD*–*trnT* (DT) with TaqI, *psaA*–*trnS*(AS) with HinfI, and *trnT*–*trnF* (TF) with AluI, following protocols described by Petit et al. ^63,103^. Primer pairs for each fragment, as described in Petit et al. (1998), were aligned to the six representative sequences and subjected to in silico restriction digestion using the Sequence Manipulation Suite ^104^ (https://www.bioinformatics.org/sms2/rest_digest.html). Fragment lengths generated from the digestion simulations were compared with Supplementary Table S3E in Nocchi et al. (2022) to assign haplotype identities. Haplotype lineage assignments were further validated by the presence of diagnostic point mutations within the *trnD*–*trnT*and *trnT*–*trnF* fragments, corresponding to restriction sites for AluI and CfoI, respectively, as described by Petit et al. (2002).

### Species differentiation

**Dataset 1** was further filtered for a minor allele frequency of 0.03 as recommended for the LEA package in R ^105^, with LD pruning repeated, and the 15 technical replicates removed. We call this **dataset 2**. We created a subset of **dataset 2** with admixed (i.e. 0.1 < q < 0.9) individuals removed and refer to this as **dataset 3**.

As a simple metric of between-species SNP differentiation, we first calculated Weir and Cockerham ^106^ per-variant pairwise F*_ST_* between the two species using **dataset 3** (i.e. the one with admixed individuals removed) in vcftools v.0.1.1 with default parameters ^90^. **Dataset 3** was used as F_st_ methods assume the data is divided into distinct subpopulations, and therefore is not designed to accommodate admixed individuals. Additionally, we used the function snmf() in R package LEA v3.17.2 ^105,107,108^ on both **dataset 2** and **dataset 3**, with arguments set to entropy = true, ploidy = 2 and K=2. This function estimates ancestry coefficients similar to those in STRUCTURE and fastSTRUCTURE, using sparse non-negative matrix factorization algorithms. Next, we used the function snmf.pvalues() in package LEA, which takes the ancestry coefficient output generated by snmf() to test for outlier loci that show excess differentiation relative to the genome-wide background. The method is similar to outlier F_ST_ approaches ^109,110^ and can be used with admixed individuals in the data ^111^, allowing for the more complete **dataset 2** to be used. Per locus p-values were calculated using the following snmf() arguments: entropy = true, K=2 and ploidy = 2. SNPs were only considered significant outliers after p-values from snmf.pvalues() were adjusted for a false discovery rate of < 0.01 using the qvalue package ^112^.

Finally, we used bedtools command intersect ^88^ in conjunction with the annotation file for the *Q. robur* genome ^64^ to identify whether the significant SNPs identified above for **dataset 2** and **dataset 3** fall within putative genes. The genes identified in this study were compared with those from other published molecular studies of species differentiation ^13,52^ and an unpublished study of the two species in mainland Europe (D. Lazic, pers.comm).

### Niche differentiation

To explore the environmental niche differentiation between *Quercus robur*, *Q. petraea*, and admixed individuals, we performed principal component analysis (PCA) separately for climate, soil, and topographic variable groups, as well as a combined PCA including all variables from 416 trees for which we had data. Climate variables comprised 19 bioclimatic variables (Bio1–Bio19) from the CHELSA-TraCE21k v.1.0 dataset ^113^(https://chelsa-climate.org/chelsa-trace21k). CHELSA-TraCE21k provides monthly climatologies for temperature and precipitation, as well as derived bioclimatic variables at 30 arcsec (∼1km) resolution in 100-year time steps for the last 21,000 years. Based on the diameter at breast height (DBH), most of our oak samples were likely established during the nineteenth and early twentieth centuries, when they were likely subjected to the strongest period of selection pressure. Consequently, we selected the average CHELSA-TraCE data from the nineteenth century as the best representation of the climatic conditions these trees experienced as saplings. Data for the sample locations were extracted using the R package terra^114^ and the longitude and latitude coordinates for each sample. Soil variables included C:N ratio, bulk density, pH, and soil moisture and were obtained from the UK Soil Observatory (UKSO) using the Country Survey dataset, which surveys the topsoil (0-15cm depth) at the 1 × 1km resolution. Topographic variables included slope, terrain ruggedness index (TRI), topographic position index (TPI), roughness, and flow accumulation and were obtained at the individual tree level from the Ordnance Survey Terrain 50 dataset (https://www.ordnancesurvey.co.uk/business-government/products/terrain-50values) at a 50m resolution with 10m height intervals and calculated in R v 4.1.1 (R Core Team, 2021) using the ‘terrain’ function in the Raster package v.3.6-20. Aspect was decomposed into northness (cosine of aspect) and eastness (sine of aspect) prior to analysis. All PCAs were performed on the correlation matrix (scale = TRUE) using the rda function from the vegan package ^115^ in R, and biplots were produced using scaling 2, which optimises the interpretation of relationships among variables. Individual trees were coloured by *Q. robur* admixture proportion to visualise the association between environmental gradients and species composition.

Because PCAs do not formally test which environmental variables predict *Q. robur* admixture proportion, we fitted a beta regression model using the betareg package^116^ in R. Originally all environmental predictors were included in the model, with *Q. robur* admixture as the response variable. A variance inflation factor implemented in the car v3.1-3 package ^117^ was performed on the full model, and the variable with the highest VIF > 10 was dropped and the model refitted. The procedure was performed until all variables had a VIF < 10. The final model contained 12 environmental variables (Bio3, Bio4, Bio8, Bio9, Bio11, Bio18, eastness, TPI, roughness, flow, C:N ratio, and pH). Model residuals were inspected for violation of assumptions, and the significance of each predictor was assessed using likelihood ratio tests (LRT).

### Growth rates

Various tree condition measures were made during the FCS. Where possible (n = 322) we calculated DBH growth for each tree from 1990 to 2019, and compared how growth rates varied with respect to ploidy (i.e. diploids vs triploids) and species, (i.e. *Q. robur*, *Q. petraea* and hybrids). To do so we ran a linear mixed model in R using the package lme4 v1.1-37 ^118^ with the difference in DBH between 1990 to 2019 as the response (cm), and ploidy and species as fixed effects. While measurement error across years may cause regression to the mean effects which need to be corrected for ^119^, DBH in 1990 and DBH in 2019 were highly correlated (r = 0.94) in our samples, making measurement corrections unwarranted. Nevertheless, we included DBH in 1990 as a covariate to control for any deviations in average growth due to differences in initial sizes or tree age. To reduce the impact of variation in environmental conditions at each site, we entered the site identity as a random effect. Due to substantial differences in sample sizes between triploids (n = 317) and diploids (n = 5) when all sites (n = 47) were considered, we also ran an alternative model including only those sites where both ploidy types were present (n = 5). This reduced the number of diploid individuals to 29. Residuals were inspected for violations of model assumptions and the response variable was log-transformed to reduce heteroscedasticity. P-values were estimated using likelihood ratio tests implemented in the drop1() function in R. Next, since environmental conditions are likely important cofactors influencing tree growth, we fitted climatic, terrain and soil variables for each site as fixed effects, rather than including site identities as random effects. In total, 35 environmental variables were included in a linear model (see the Niche differentiation section above), with species, ploidy and DBH in 1990 as covariates. A variance inflation factor using the package car v3.1-3 ^117^ was performed to investigate collinearity between variables following the procedure delineated above. P-values of the reduced model were estimated using F tests.

## References

1. Greville, R. K. VIII.On the botanical characters of the British oaks. Trans. - Bot. Soc. Edinb. (1971) 1, 65–69 (1844).

2. Darwin, C. On the Origin of Species by Means of Natural Selection, Or, The Preservation of Favoured Races in the Struggle for Life. (John Murray, London :, 1859).

3. Cousens, J. E. Variation of some diagnostic characters of the sessile and pedunculate oaks and their hybrids in Scotland. Watsonia 5, 273–286 (1963).

4. Cousens, J. The status of the pedunculate and sessile oaks in Britain. Watsonia 6, 161–176 (1965).

5. Moss, C. E. British Oaks. J. Bot. 48, 1–8, 33–39 (1910).

6. Gardiner, A. S. Pedunculate and Sessile Oak (Quercus robur L. and Quercus petraea (Mattuschka) Liebl.): A Review of the Hybrid Controversy. Forestry (Lond.) 43, 151–160 (1970).

7. Gardiner, A. S. A history of the taxonomy and distribution of the native oak species. in The British Oak (eds. Morris, M. G. & Perring, F. H.) 3–26 (The Botanical Society of the British Isles, Faringdon, Berkshire, UK, 1974).

8. Rushton, B. Quercus robur L. and Quercus petraea (Matt.) Liebl.: a multivariate approach to the hybrid problem. 2. The geographical distribution of population types. New Journal of Botany 12, 209–224 (1979).

9. Rushton, B. S. Natural hybridization within the genus Quercus L. Ann. Sci. For. 50, 73s–90s (1993).

10. Rushton, B. An analysis of variation of leaf characters in Quercus robur L. and Quercus petraea (Matt.) Liebl. population samples from Northern Ireland. Irish Forestry (1983).

11. Cottrell, J. E. et al. Distribution of chloroplast DNA variation in British oaks (Quercus robur and Q. petraea): the influence of postglacial colonisation and human management. For. Ecol. Manage. 156, 181–195 (2002).

12. Gerber, S. et al. High rates of gene flow by pollen and seed in oak populations across Europe. PLoS One 9, e85130 (2014).

13. Nocchi, G. et al. Genomic structure and diversity of oak populations in British parklands. Plants People Planet 4, 167–181 (2022).

14. Kremer, A. et al. Leaf morphological differentiation between Quercus robur and Quercus petraea is stable across western European mixed oak stands. Ann. For. Sci. 59, 777–787 (2002).

15. Petit, R. J., Bodénès, C., Ducousso, A., Roussel, G. & Kremer, A. Hybridization as a mechanism of invasion in oaks: Research review. New Phytol. 161, 151–164 (2004).

16. Rellstab, C. et al. Signatures of local adaptation in candidate genes of oaks (Quercusspp.) with respect to present and future climatic conditions. Mol. Ecol. 25, 5907–5924 (2016).

17. Viscosi, V., Lepais, O., Gerber, S. & Fortini, P. Leaf morphological analyses in four European oak species (Quercus) and their hybrids: A comparison of traditional and geometric morphometric methods. Plant Biosyst. 143, 564–574 (2009).

18. Aas, G. Taxonomical impact of morphological variation in Quercus robur and Q petraea: a contribution to the hybrid controversy. Ann. Sci. For. 50, 107s–113s (1993).

19. Kremer, A. & Hipp, A. L. Oaks: an evolutionary success story. New Phytol. 226, 987–1011 (2020).

20. Leroy, T. et al. Extensive recent secondary contacts between four European white oak species. New Phytol. 214, 865–878 (2017).

21. Petit, R. J. et al. Identification of refugia and post-glacial colonisation routes of European white oaks based on chloroplast DNA and fossil pollen evidence. For. Ecol. Manage. 156, 49–74 (2002).

22. Ferris, C., Oliver, R. P., Davy, A. J. & Hewitt, G. M. Native oak chloroplasts reveal an ancient divide across Europe. Mol. Ecol. 2, 337–344 (1993).

23. Roloff, A. & Bärtels, A. Flora Der Gehölze. (Ulmer, Stuttgart, 2008).

24. Cuypers, V. & Reydon, T. A. C. An oak is an oak, or not? Understanding and dealing with confusion and disagreement in biological classification. Biol. Philos. 38, (2023).

25. Goodwin, H. & Flandrian, J. D. History of oak in the British Isles. in The British Oak (eds. Morris, M. G. & Perring, F. H.) 62–79 (The Botanical Society of the British Isles, Faringdon, Berkshire, UK, 1974).

26. Lowe, A. et al. Route, speed and mode of oak postglacial colonisation across the British Isles: Integrating molecular ecology, palaeoecology and modelling approaches. Bot. J. Scotl. 57, 59–81 (2005).

27. POWO. Plants of the World Online. Plants of the World Online. Facilitated by the Royal Botanic Gardens, Kew. https://powo.science.kew.org/ (2026).

28. Bordacs, S. & Ducousso, A. EUFORGEN Technical Guidelines for Genetic Conservation and Use for Pedunculate and Sessile Oaks (Quercus Robur and Q. Petraea). (2004).

29. Stroh, P. A., Walker, K. J., Humphrey, T. A., Pescott, O. L. & Burkmar, R. J. Plant Atlas 2020: Mapping Changes in the Distribution of the British and Irish Flora. (Princeton University Press, Princeton, NJ, 2023).

30. Jones, E. W. Quercus L. J. Ecol. 47, 169 (1959).

31. Stace, C. A. New Flora of the British Isles, Edition 4. (C&M Floristics, Stowmarket, England, 2019).

32. Maxwell, H. Trees: A Woodland Notebook. (James Maclehose and Sons, Glasgow, 1915).

33. Eaton, E, Caudullo, G, Oliveira, S. & De, R. D. *Quercus robur* and *Quercus petraea* in Europe: distribution, habitat, usage and threats. in European atlas of forest tree species (ed. de Rigo D. Caudullo G. Houston Durrant T. Mauri A, S.-M.-A. J.) 160–163 (Publ. Off. EU, Luxembourg, 2016).

34. Rackham, O. Woodlands. (William Collins, London, England, 2015).

35. Savill, P. The Silviculture of Trees Used in British Forestry. (cabidigitallibrary.org, 2013).

36. Rackham, O. Trees and Woodland in the British Landscape. (Hachette, UK, 2020).

37. Cruickshank, T. The Practical Planter. (William Blackwood, Edinburgh, 1830).

38. Konatowska, M., Młynarczyk, A., Rutkowski, P. & Kujawa, K. Impact of Site Conditions on Quercus robur and Quercus petraea Growth and Distribution Under Global Climate Change. Remote Sens. (Basel) 16, 4094 (2024).

39. Ponton, S. et al. Carbon isotope discrimination and wood anatomy variations in mixed stands of *Quercus robur* and *Quercus petraea*. Plant Cell Environ. 24, 861–868 (2001).

40. Nicolescu, V.-N. et al. Management of sessile oak (Quercus petraea (Matt.) Liebl.), a major forest species in Europe. J. For. Res. 36, (2025).

41. Diaz-Maroto, I. J., Vila-Lameiro, P., Ramón-Pumar, M., Alañón, E. & Diaz-Maroto, M. C. The different occurrence conditions ofQuercus roburL. andQuercus petraea(Mattuschka) Liebl. on current habitat in Galicia, NW Iberian Peninsula. Scand. J. For. Res. 30, 122–134 (2015).

42. Cochard, H., Bréda, N., Granier, A. & Aussenac, G. Vulnerability to air embolism of three European oak species (Quercus petraea (Matt) Liebl, Q pubescens Willd, Q robur L). Ann. Sci. For. 49, 225–233 (1992).

43. Hauck, M. et al. Heat tolerance of temperate tree species from Central Europe. For. Ecol. Manage. 580, 122541 (2025).

44. Epron, D. & Dreyer, E. Long-term effects of drought on photosynthesis of adult oak trees [Quercus petraea (Matt.) Liebl. and Quercus robur L.] in a natural stand. New Phytol. 125, 381–389 (1993).

45. Schmull, M. & Thomas, F. M. Morphological and physiological reactions of young deciduous trees (Quercus robur L., Q. petraea [Matt.] Liebl., Fagus sylvatica L.) to waterlogging. Plant Soil 225, 227–242 (2000).

46. Wagner, P. A. & Dreyer, E. Interactive effects of waterlogging and irradiance on the photosynthetic performance of seedlings from three oak species displaying different sensitivities (Quercus robur, Q petraea and Q rubra). Ann. Sci. For. 54, 409–429 (1997).

47. Lazic, D., Hipp, A. L., Carlson, J. E. & Gailing, O. Use of genomic resources to assess adaptive divergence and introgression in oaks. Forests 12, 690 (2021).

48. Reutimann, O., Gugerli, F. & Rellstab, C. A species-discriminatory single-nucleotide polymorphism set reveals maintenance of species integrity in hybridizing European white oaks (Quercus spp.) despite high levels of admixture. Ann. Bot. 125, 663–676 (2020).

49. Leroy, T. et al. Massive postglacial gene flow between European white oaks uncovered genes underlying species barriers. New Phytol. 226, 1183–1197 (2020).

50. Guichoux, E. et al. Outlier loci highlight the direction of introgression in oaks. Mol. Ecol. 22, 450–462 (2013).

51. Saintagne, C. et al. Distribution of genomic regions differentiating oak species assessed by QTL detection. Heredity (Edinb.) 92, 20–30 (2004).

52. Leroy, T. et al. Adaptive introgression as a driver of local adaptation to climate in European white oaks. New Phytol. 226, 1171–1182 (2020).

53. Lepais, O., Roussel, G., Hubert, F., Kremer, A. & Gerber, S. Strength and variability of postmating reproductive isolating barriers between four European white oak species. Tree Genet. Genomes 9, 841–853 (2013).

54. Steinhoff, S. Controlled crosses between pendunculate and sessile oak - results and conclusion. Allg. Forst- Jagdztg. 169, 163–168 (1998).

55. Chybicki, I. J. & Burczyk, J. Seeing the forest through the trees: comprehensive inference on individual mating patterns in a mixed stand of Quercus robur and Q. petraea. Ann. Bot. 112, 561–574 (2013).

56. Lagache, L., Klein, E. K., Ducousso, A. & Petit, R. J. Distinct male reproductive strategies in two closely related oak species. Mol. Ecol. 23, 4331–4343 (2014).

57. Truffaut, L. et al. Fine-scale species distribution changes in a mixed oak stand over two successive generations. New Phytol. 215, 126–139 (2017).

58. Curtu, A. L., Gailing, O. & Finkeldey, R. Evidence for hybridization and introgression within a species-rich oak (Quercus spp.) community. BMC Evol. Biol. 7, 218 (2007).

59. Mitchell, R. J. et al. Collapsing foundations: The ecology of the British oak, implications of its decline and mitigation options. Biol. Conserv. 233, 316–327 (2019).

60. Great Britain: Forestry Commission. Analysis of the Changes in Forest Condition in Britain, 1989-92. (Stationery Office Books, Norwich, England, 1995).

61. Great Britain: Forestry Commission. Monitoring of Forest Conditions in the United Kingdom 1988. (Stationery Office Books, Norwich, England, 1989).

62. Innes, J. L. & Boswell, R. C. Forest Health Surveys 1987 Part 1: Results. (1987).

63. Petit, R. J. et al. Chloroplast DNA variation in European white oaks. Phylogeography and patterns of diversity based on data from over 2600 populations. [Erratum: Mar 17, 2003, v. 176 (1/3), p. 595-599.]. For. Ecol. Manage. 156, 5–26 (2002).

64. Plomion, C. et al. Oak genome reveals facets of long lifespan. Nat. Plants 4, 440–452 (2018).

65. Lepais, O. et al. Species relative abundance and direction of introgression in oaks. Mol. Ecol. 18, 2228–2242 (2009).

66. Lang, T., et al. High-quality SNPs from genic regions highlight introgression patterns among European white oaks (Quercus petraea and Q. robur). bioRxiv 388447 (2021).

67. Degen, B., Blanc-Jolivet, C., Mader, M., Yanbaeva, V. & Yanbaev, Y. Introgression as an important driver of geographic genetic differentiation within European White Oaks. Forests 14, 2279 (2023).

68. Belton, S., Fox, E., Degen, B. & Kelleher, C. T. Extensive hybridisation and introgression between two European white oaks at the western limit of their range. Conserv. Genet. 27, (2026).

69. Carleial, R. et al. Genomic basis for resistance to acute oak decline and mildew infection in English oak. bioRxiv 2026.03.04.709495 (2026).

70. Eriksson, G. & Jonsson, A. A review of the genetics ofBetula. Scand. J. For. Res. 1, 421–434 (1986).

71. Bjerkan, K. N. et al. Genetic variation and temperature affects hybrid barriers during interspecific hybridization. Plant J. 101, 122–140 (2020).

72. Muir, G. & Schlötterer, C. Evidence for shared ancestral polymorphism rather than recurrent gene flow at microsatellite loci differentiating two hybridizing oaks (*Quercus* spp.): SHARED ANCESTRAL VARIATION IN OAKS. Mol. Ecol. 14, 549–561 (2005).

73. Hubert, J. Selecting the Right Provenance of Oak for Planting in Britain. https://www.cabidigitallibrary.org/doi/full/10.5555/20063048227 (2005) doi:10.5555/20063048227.

74. Lévy, G., Becker, M. & Duhamel, D. A comparison of the ecology of pedunculate and sessile oaks: Radial growth in the centre and northwest of France. For. Ecol. Manage. 55, 51–63 (1992).

75. Buriánek, V., Benedíková, M. & Kyseláková, J. Evaluation of twenty-years-old pedunculate and sessile oak provenance trial. J. For. Sci. 57, 153–169 (2011).

76. Nyamjav, B., Herzog, S. & Krabel, D. Reaction of oak seedlings (Quercus robur L. and Quercus petraea (Matt.) Liebl.) to water limitation. Austrian J. For. Sci. 139, 117–136 (2022).

77. Butorina, A. K. Cytogenetic study of diploid and spontaneous triploid oaks, Quercus robur L. Ann. Sci. For. 50, 144s–150s (1993).

78. Dzialuk, A., Chybicki, I., Welc, M., Sliwinska, E. & Burczyk, J. Presence of triploids among oak species. Ann. Bot. 99, 959–964 (2007).

79. Naujoks, G., Hertel, H. & Ewald, D. Characterization and propagation of an adult triploid pedunculate oak (Quercus robur L.). Silvae Genetica 44, 282–286 (1995).

80. Mather, R. A., Freer-Smith, P. H. & Savill., P. S. Analysis of the Changes in Forest Condition in Britain, 1989 to 1992. (1995).

81. Brown, N., Jeger, M., Kirk, S., Xu, X. & Denman, S. Spatial and temporal patterns in symptom expression within eight woodlands affected by Acute Oak Decline. For. Ecol. Manage. 360, 97–109 (2016).

82. Chen, S., Zhou, Y., Chen, Y. & Gu, J. fastp: an ultra-fast all-in-one FASTQ preprocessor. Bioinformatics 34, i884–i890 (2018).

83. Bolger, A. M., Lohse, M. & Usadel, B. Trimmomatic: a flexible trimmer for Illumina sequence data. Bioinformatics 30, 2114–2120 (2014).

84. Li, H. Aligning sequence reads, clone sequences and assembly contigs with BWA-MEM. arXiv [q-bio.GN] (2013).

85. Danecek, P. et al. Twelve years of SAMtools and BCFtools. Gigascience 10, giab008 (2021).

86. McKenna, A. et al. The Genome Analysis Toolkit: a MapReduce framework for analyzing next-generation DNA sequencing data. Genome Res. 20, 1297–1303 (2010).

87. Poplin, R., et al. Scaling accurate genetic variant discovery to tens of thousands of samples. bioRxiv 201178 (2018) doi:10.1101/201178.

88. Quinlan, A. R. & Hall, I. M. BEDTools: a flexible suite of utilities for comparing genomic features. Bioinformatics 26, 841–842 (2010).

89. R Foundation for Statistical Computing, Vienna, Austria. R: A Language and Environment for Statistical Computing. (2022).

90. Danecek, P. et al. The variant call format and VCFtools. Bioinformatics 27, 2156–2158 (2011).

91. Zohren, J. et al. Unidirectional diploid-tetraploid introgression among British birch trees with shifting ranges shown by restriction site-associated markers. Mol. Ecol. 25, 2413–2426 (2016).

92. Fletcher, K., Han, R., Smilde, D. & Michelmore, R. Variance of allele balance calculated from low coverage sequencing data infers departure from a diploid state. BMC Bioinformatics 23, 150 (2022).

93. Raj, A., Stephens, M. & Pritchard, J. K. fastSTRUCTURE: variational inference of population structure in large SNP data sets. Genetics 197, 573–589 (2014).

94. Purcell, S. et al. PLINK: a tool set for whole-genome association and population-based linkage analyses. Am. J. Hum. Genet. 81, 559–575 (2007).

95. Neophytou, C. Bayesian clustering analyses for genetic assignment and study of hybridization in oaks: effects of asymmetric phylogenies and asymmetric sampling schemes. Tree Genet. Genomes 10, 273–285 (2014).

96. Lemaire, J. Le chêne autrement: Produire du chêne de qualité en moins de 100 ans en futaie régulière. (2010).

97. Gathercole, L. A. P. et al. Evidence for the Widespread Occurrence of Bacteria Implicated in Acute Oak Decline from Incidental Genetic Sampling. For. Trees Livelihoods 12, 1683 (2021).

98. Wickham, H., Navarro, D. & Pedersen, T. L. ggplot2: Elegant Graphics for Data Analysis (3e). https://ggplot2-book.org/.

99. Dierckxsens, N., Mardulyn, P. & Smits, G. NOVOPlasty: de novo assembly of organelle genomes from whole genome data. Nucleic Acids Res. 45, e18 (2017).

100. Katoh, K. & Standley, D. M. MAFFT multiple sequence alignment software version 7: improvements in performance and usability. Mol. Biol. Evol. 30, 772–780 (2013).

101. Capella-Gutiérrez, S., Silla-Martínez, J. M. & Gabaldón, T. trimAl: a tool for automated alignment trimming in large-scale phylogenetic analyses. Bioinformatics 25, 1972–1973 (2009).

102. Leigh, J. W. & Bryant, D. Popart: Fulllfeature software for haplotype network construction. Methods Ecol. Evol. 6, 1110–1116 (2015).

103. Petit, R. J., Demesure, B. & Dumolin, S. cpDNA and mtDNA Primers in Plants. in Molecular Tools for Screening Biodiversity 256–261 (Springer Netherlands, Dordrecht, 1998).

104. Stothard, P. The sequence manipulation suite: JavaScript programs for analyzing and formatting protein and DNA sequences. Biotechniques 28, 1102, 1104 (2000).

105. Frichot, E. & François, O. LEA: An R package for landscape and ecological association studies. Methods in Ecology and Evolution 6, 925–929 (2015).

106. Weir, B. S. & Cockerham, C. C. Estimating F-statistics for the analysis of population structure. Evolution 38, 1358 (1984).

107. Gain, C. & François, O. LEA 3: Factor models in population genetics and ecological genomics with R. Mol. Ecol. Resour. 21, 2738–2748 (2021).

108. Jumentier, B., Caye, K., Heude, B., Lepeule, J. & François, O. Sparse latent factor regression models for genome-wide and epigenome-wide association studies. Stat. Appl. Genet. Mol. Biol. 21, (2022).

109. Beaumont, M. A. & Nichols, R. A. Evaluating Loci for Use in the Genetic Analysis of Population Structure. Proceedings of the Royal Society of London B: Biological Sciences 263, 1619–1626 (1996).

110. Lotterhos, K. E. & Whitlock, M. C. Evaluation of demographic history and neutral parameterization on the performance of FST outlier tests. Mol. Ecol. 23, 2178–2192 (2014).

111. Martins, H., Caye, K., Luu, K., Blum, M. G. B. & François, O. Identifying outlier loci in admixed and in continuous populations using ancestral population differentiation statistics. bioRxiv (2016) doi:10.1101/054585.

112. jdstorey/qvalue: Q-value estimation for false discovery rate control. https://rdrr.io/github/jdstorey/qvalue/ (2023).

113. Karger, D. N., Wilson, A. M., Mahony, C., Zimmermann, N. E. & Jetz, W. Global daily 1 km land surface precipitation based on cloud cover-informed downscaling. Sci. Data 8, 307 (2021).

114. Hijmans, R. J. terra: Spatial Data Analysis. CRAN: Contributed Packages The R Foundation 10.32614/cran.package.terra (2020).

115. Oksanen, J., et al. Community Ecology Package. (CRAN, 2025).

116. Cribari-Neto, F. & Zeileis, A. Beta Regression inR. J. Stat. Softw. 34, 1–24 (2010).

117. Fox, J. & Weisberg, S. An R Companion to Applied Regression. (SAGE Publications, Thousand Oaks, CA, 2018).

118. Bates, D., Mächler, M., Bolker, B. & Walker, S. Fitting linear mixed-effects models Usinglme4. J. Stat. Softw. 67, (2015).

119. Kelly, C. & Price, T. D. Correcting for regression to the mean in behavior and ecology. Am. Nat. 166, 700–707 (2005).

