## Supplementary Material for "Genomic differentiation of British oaks"

**SUPPLEMENTARY FIGURES**

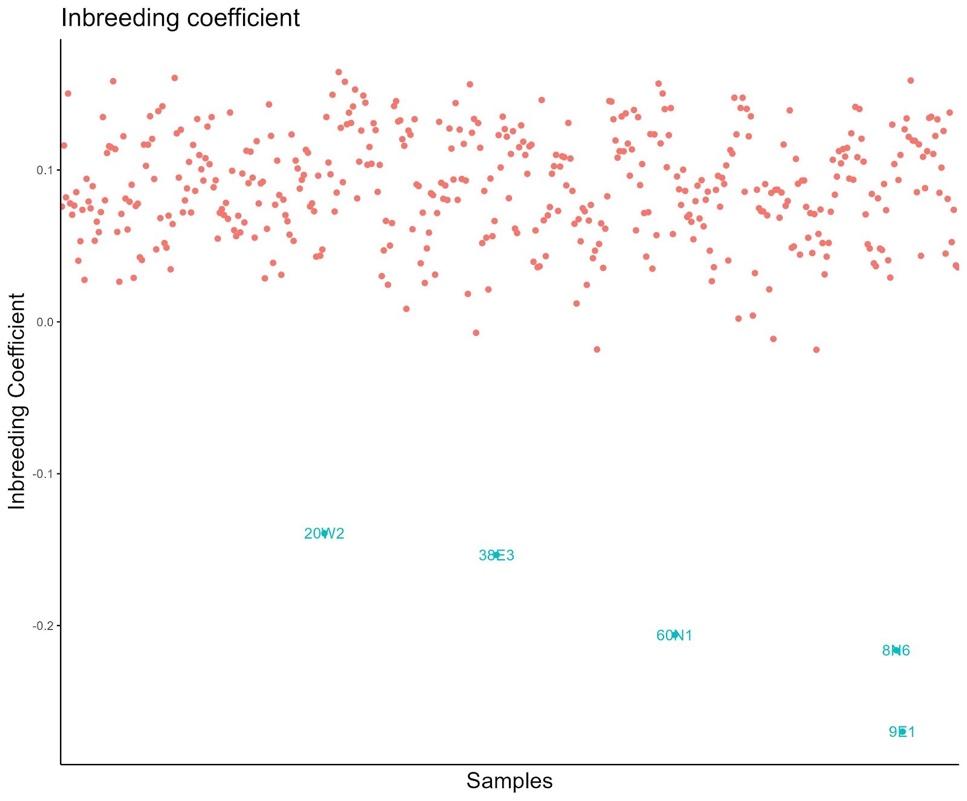

**Supplementary Figure 1:** Inbreeding coefficient per sample. Suspected triploids show pronounced negative inbreeding coefficients indicating excess heterozygosity, and can be seen in the lower part of the plot, coloured blue.

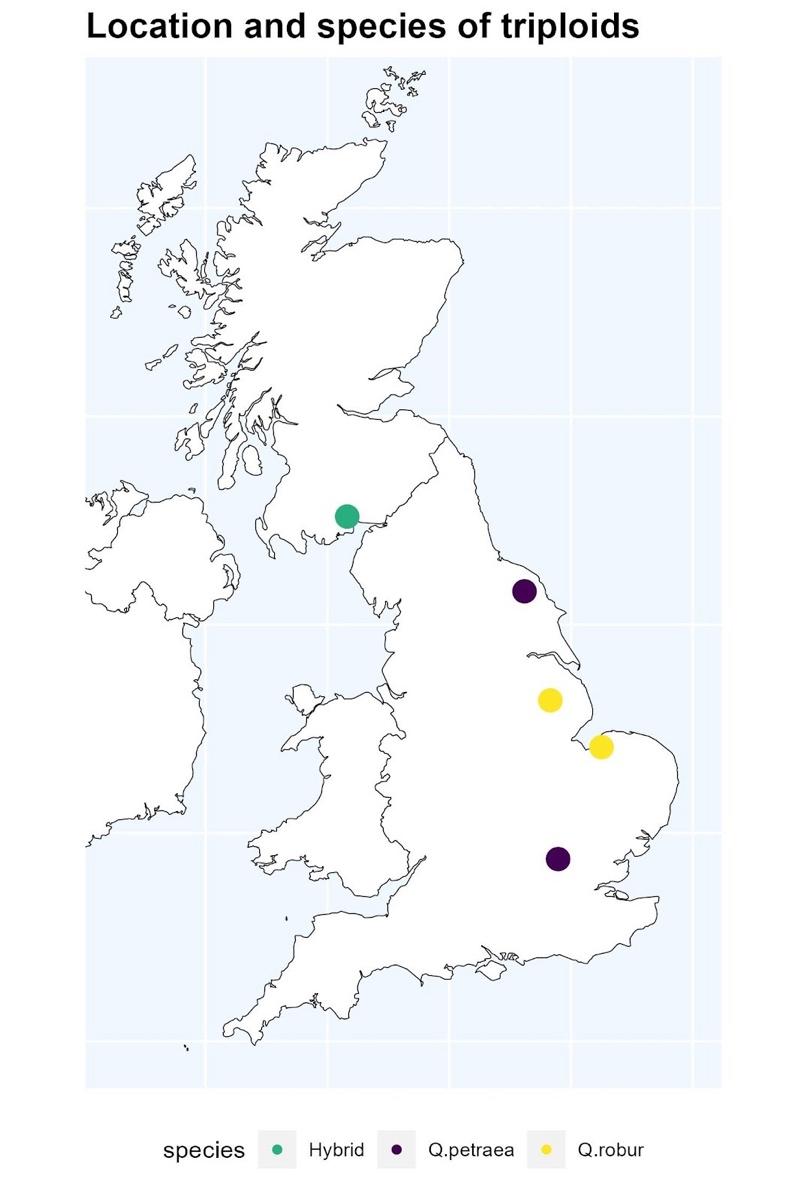

**Supplementary Figure 2**: Location and species allocations of suspected triploid individuals. Two are

*Q. robur*, two are *Q. petraea* and one is a back-crossed *Q. petraea*

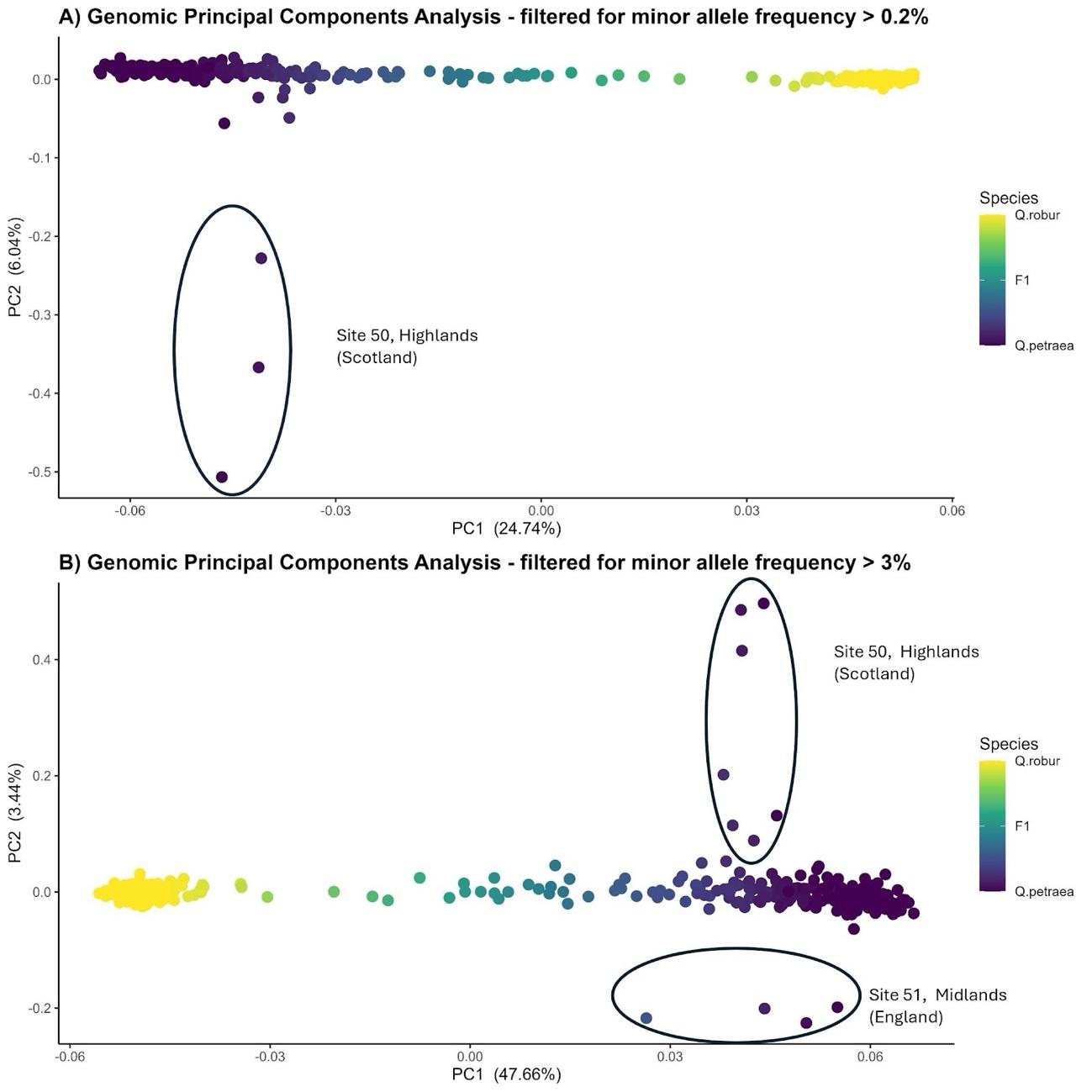

**Supplementary Figure 3:** genetic principal components analysis summarising genetic variation in multiple populations of British oak. PC1, accounting for 48% of the variation, separates the two native species (*Q. robur* and *Q. petraea*), with hybrids/admixed samples lying on a continuum between the species clusters. PC2, accounting for 3% of the variation, separates out samples from two sites (50 & 51) which are highlighted with ellipses. Data filtered for minor allele frequency of (a) >0.2% and (b) > 3%.

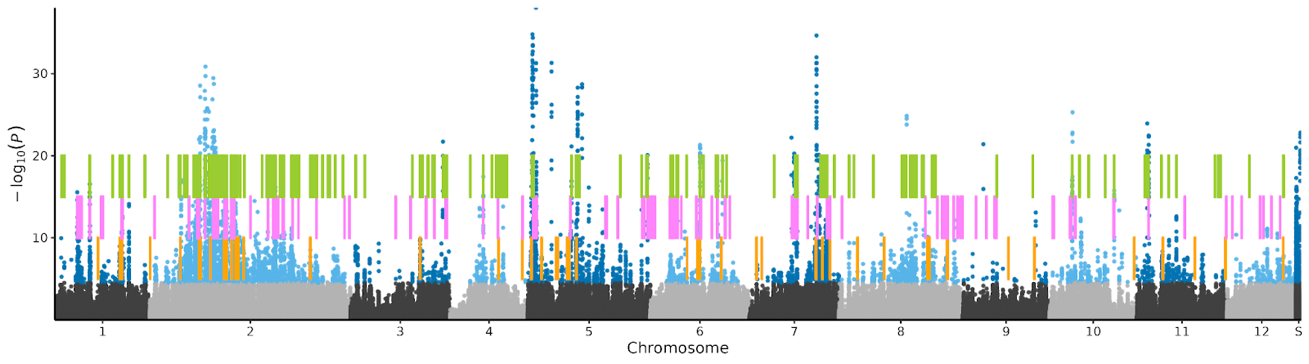

**Supplementary Figure 4:** Manhattan plot of p-values from tests for outlier loci showing excess differentiation between *Q. robur* and *Q. petraea* relative to the genome-wide background based on Dataset 2 (which includes admixed individuals) using snmf. SNPs with a q value FDR <0.01 are highlighted in blue. Coloured vertical lines show locations of genes identified as differentiated between the two species in previous studies: Green - Lazic̀ et al. (pers. comm); Pink: Leroy et al. (Leroy, Louvet, et al. 2020); Yellow - Nocchi et al. (Nocchi et al. 2022).

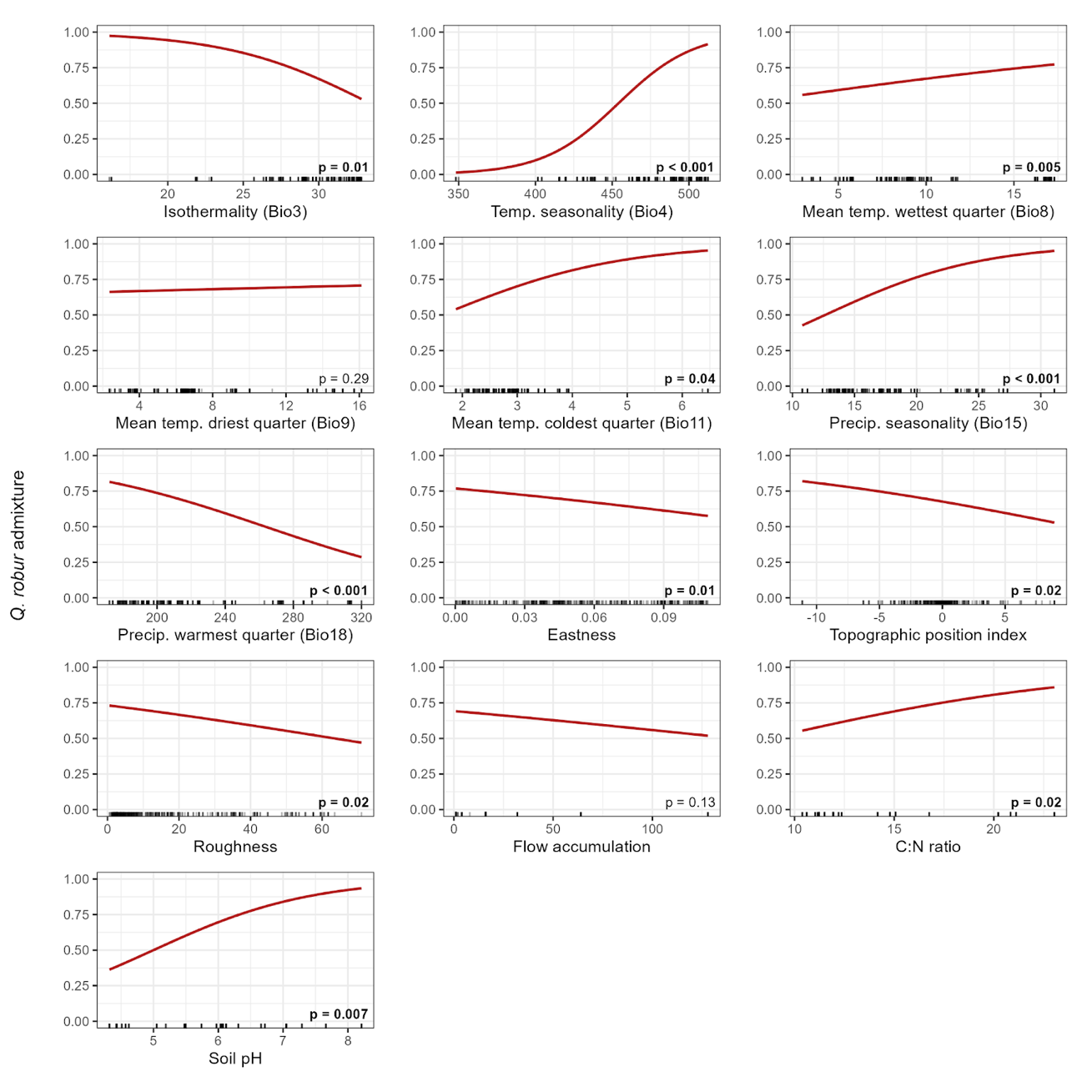

**Supplementary Figure 5:** Marginal effects of a beta regression model testing for associations between environmental predictors and the proportion of *Q. robur* admixture (n = 416 trees). Tick marks on the x-axis represent the observed distribution of each variable, and the red lines are the regression slopes. Likelihood ratio test p-values are shown in each panel; bold values indicate statistical significance (p < 0.05). The response was modelled on the logit scale with a logit link function; predicted values were back-transformed to the proportion scale for display.
